# Most Upf1-associated mRNAs have short poly(A) tails, lack a premature termination codon and are targeted by NMD

**DOI:** 10.1101/2025.02.21.639497

**Authors:** Toni Gouhier, Cosmin Saveanu, Gwenael Badis

## Abstract

Nonsense-mediated mRNA decay (NMD) is a conserved eukaryotic surveillance pathway known to degrade mRNAs containing premature termination codons (PTCs). A distance long enough between the stop codon and the poly(A)-binding protein (Pab1) is required for mRNA recognition by the NMD factors Upf1, Upf2 and Upf3. Using Nanopore direct RNA sequencing, we show that PTC-containing NMD targets account for only 6% of Upf1-associated RNA and have long poly(A) tails, indicating that Upf1-binding occurs prior to RNA deadenylation. Conversely, most Upf1-associated mRNAs have short poly(A) tails, lack a PTC and correspond to highly expressed genes. A short poly(A) tail is thus an important feature of NMD targets, redefining the scope of this RNA degradation pathway. We propose a model in which loss of Pab1-binding to short poly(A)-tailed mRNAs impairs translation termination and dictates the recruitment of the NMD machinery, uncovering a hitherto unknown role of NMD in the degradation of these transcripts.

## INTRODUCTION

Nonsense-mediated mRNA decay (NMD) is a major cytoplasmic surveillance pathway, highly conserved across eukaryotes, that eliminates aberrant transcripts containing a premature termination codon (PTC) to prevent the production of truncated, potentially deleterious proteins. Upstream frameshift 1 (Upf1), the NMD core factor, is an ATP-dependent RNA helicase. In yeast, Upf1 associates with at least two distinct subcomplexes: the Upf1-23 complex, comprising Upf1, Upf2, and Upf3, and the Upf1-decapping complex, which includes the decapping enzymes Dcp1/2, Edc3, Ebs1 and Nmd4 (Dehecq *et al*, 2018). The Upf1-23 complex plays a central role in NMD activation and interacts with the translation release factors eRF1 and eRF3, which are crucial for efficient ribosome release (Czaplinski *et al*, 1998; Wang *et al*, 2001). In contrast, the Upf1-decapping complex mediates mRNA decapping, triggering subsequent 5′ to 3′ exonucleolytic degradation by Xrn1. Remarkably, this process occurs independently of prior deadenylation (Muhlrad & Parker, 1994, 1999; Cao & Parker, 2003). According to the faux 3’UTR consensus model, NMD-targeted mRNAs experience defective translation termination due to the presence of a PTC or an unusually long 3’UTR (Kervestin & Jacobson, 2012; Hug *et al*, 2016; Losson & Lacroute, 1979). This model posits that inefficient translation termination arise from the absence of the stimulatory role of a proximal Pab1 on eRF3 (Celik *et al*, 2015).

In contrast, for “normal” non-PTC-containing mRNAs (noPTC), translation termination is facilitated by the interaction between eRF3 and the poly(A)-binding protein Pab1, recruited close to the stop codon by the proximal poly(A) tail (Roque *et al*, 2015). In addition, noPTC general mRNA decay predominantly relies on a deadenylation-dependent mechanism, where deadenylation serves as a prerequisite for subsequent decapping and degradation (Beelman & Parker, 1995; Chen & Shyu, 2011). Deadenylation is mostly carried out by two enzymatic complexes, Pan2/Pan3 and CCR4/NOT, both contributing to the shortening of the poly(A) tail to a residual length of 10–15 nucleotides. The shortened poly(A) tail is then recognized by the Lsm1-7/Pat1 complex, which facilitates decapping through the recruitment of Dcp1/2. Once decapped, mRNAs are predominantly degraded from 5′ to 3′ by the Xrn1 exonuclease, or in a minor way, from 3′ to 5′ by the Ski7-exosome complex (see Coller & Parker, 2004 for a review).

PTC-containing mRNAs are common and include mRNA with upstream open reading frames, retained introns, or frameshift mutations, as well as non-coding RNAs like XUTs and SUTs (see Nickless *et al*, 2017 for a review). Initially thought to be rare, transcripts with alternative transcription start sites that generate PTCs have been identified in more than half of yeast genes. These transcripts, although representing a minority of the transcriptome, are selectively targeted and degraded by NMD (Malabat *et al*, 2015). Notably, NMD also downregulates a substantial fraction of non-mutated mRNAs lacking canonical PTC features (He *et al*, 2003; Guan *et al*, 2006).

Several strategies have been employed to assess the transcriptome-wide scope of mRNA degradation by NMD. An initial approach involved the identification of transcripts that were up-regulated or stabilized upon Upf1 inactivation (He *et al*, 2003; Celik *et al*, 2017; Tani *et al*, 2012). These studies revealed the most efficiently degraded NMD substrates, which presumably correspond to homogeneous populations of PTC-containing mRNAs, estimated to represent no more than 5% of all mRNAs (He *et al*, 2003). To identify Upf1 direct targets, RNA immunoprecipitation followed by sequencing (RIP-seq) and cross-linking-based methods led to a different outcome. These approaches identified Upf1 binding not only to PTC-containing mRNAs but also to a wide range of noPTC transcripts, suggesting a broader role for Upf1 in general mRNA decay (Hurt *et al*, 2013; Zünd *et al*, 2013; Kurosaki *et al*, 2018).

The mentioned previous studies did not measure the poly(A) tails of individual Upf1-associated RNA. Each transcript is a complex mix of RNAs at different stages of polyadenylation, in relation with their degradation rates. Recent studies established that deadenylation is not required for decapping and degradation of most yeast mRNAs under normal growth conditions (Czarnocka-Cieciura *et al*, 2024; Audebert *et al*, 2024), or during meiosis (Wiener *et al*, 2021). Consistently, unstable mRNAs, which are rapidly decapped and degraded 5’ to 3’, tend to have long poly(A) tails, whereas stable mRNAs harbor shorter poly(A) tails (Subtelny *et al*, 2014; Tudek *et al*, 2021). At steady state, mRNAs expressed from a single gene is a mix of molecules with long (>50 to 80 adenosine residues in yeast, up to 300 in humans), intermediate (25–50A), short (<25A), or even lacking poly(A) tails. Transcripts can be capped or decapped and may undergo partial 5’ degradation. Since NMD substrates are unstable, they are degraded before having time to undergo deadenylation (Audebert *et al*, 2024). Thus, the status of the poly(A) tail in transcript populations provides important, and previously unexplored information about their degradation and targeting by NMD.

On one hand, NMD is deadenylation-independent and targets mRNA with long poly(A) tails (Audebert *et al*, 2024). On the other hand, the Lsm1-7/Pat1 complex is involved in the recognition of short polyadenylated mRNA, facilitating decapping (Tharun & Parker, 2001). Surprisingly, Upf1, Lsm1, Pat1 and Xrn1 degradation factors can coexist in the same mRNP (Dehecq *et al*, 2018), Since the recognition of short polyadenylated mRNAs by the Lsm1-7/Pat1 complex enhances decapping (Tharun & Parker, 2001), these biochemical data suggest that Upf1 may also play a role in the degradation of short poly(A)-tailed mRNAs as a general decay factor beyond its established role in NMD.

Despite extensive efforts to understand NMD mechanisms, the rules governing transcript recognition by Upf1 and the NMD machinery remain unclear. This is particularly true for mRNA that do not contain a PTC but are still associated with Upf1. To address this point, here we used a combination of high-throughput RNA and RIP sequencing in *Saccharomyces cerevisiæ*, to show that Upf1 associates not only with PTC-containing NMD substrates with long poly(A) tails, but also with a substantial fraction of transcripts lacking a PTC and carrying short poly(A) tails. This RNA subpopulation accumulates in heavy polysome fractions and is stabilized in the absence of a functional NMD machinery. Our results show that this subfraction of short poly(A)-tailed mRNAs, originating from stable and highly translated mRNA, undergoes NMD-dependent degradation and consists of stable and highly translated mRNAs, revealing a novel role for the NMD machinery in the degradation of mRNA beyond PTC-containing transcripts.

## RESULTS

### A subpopulation of mRNAs lacking PTCs is associated to Upf1

Because transcript binding to Upf1 is the first step in NMD activation, Upf1-TAP associated RNAs were purified and sequenced by Illumina (RIPseq) (Figure 1A). RIPseq were performed under conditions that have been previously used to describe the architecture of yeast NMD complexes (Dehecq *et al*, 2018). We found a robust association of known NMD-targeted transcripts (**XUT/SUT** and **PTC**-containing mRNAs) with Upf1 (blue dots in Figure 1B). We distinguished non-PTC containing transcripts according to their enrichment in Upf1 immunoprecipitates (**no_PTC_enr** red dots and **noPTC**, pink dots). Abundant mitochondrial-encoded mRNAs (yellow) were hardly detected, indicating a low immunoprecipitation background.

**Figure 1.**
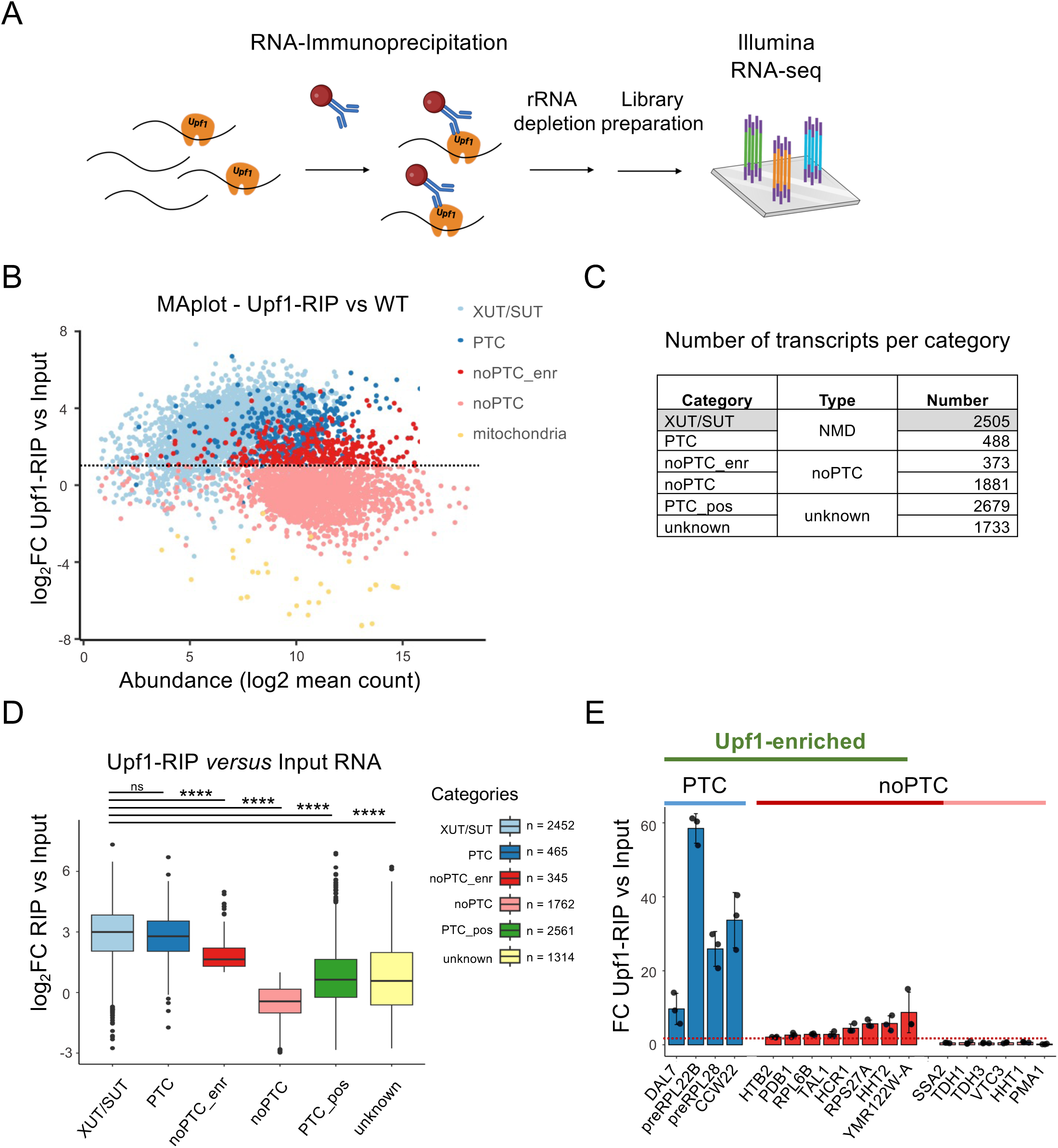
A subpopulation of mRNAs lacking PTCs is associated to Upf1. (A) Schematic representation of RNA-Immunoprecipitation (RIP) followed by Illumina sequencing. (B) MAplot representing the log2FC between Upf1-RIP and WT as a function of abundance (A = log2 mean count in Upf1-RIP and WT). Each transcript type is represented as follow: XUT/SUT (light blue), PTC (dark blue), noPTC-Upf1enr (red), noPTC (pink), and mitochondrial transcripts as a control (yellow). Dotted lines indicate a 2-fold threshold (log2FC = 1). NoPTC_enr were defined as noPTC transcripts enriched more than two-fold in Upf1 RIP (log2FC Upf1 RIP vs input>1). Three biological replicates of each condition were used for all the experiments of this study. (C) Number of transcripts of each type in the different categories. Grey-shaded boxes represent XUT/SUT transcripts, and white boxes detail protein-coding gene sub-categories (PTC, noPTC, PTC_pos, and unknown). (D) Boxplots comparing Upf1-RIP vs Input log2FC for mentioned categories of transcripts. The cartoon describes categories (as in (A) and corresponding number of detected transcripts. Pvalues (Tukey HSD test): ns = p> 0.05, * = p<0,05, ** = p<0,01, *** = p<0,001 and **** = p<0,0001. See also Table S2 for detailed pvalues and Table S3 for normalized count. (E) Barplot comparing Upf1-RIP versus Input fold change (FC) for PTC and noPTC examples.

As a first step in understanding the rules that govern transcript association with Upf1, we classified annotated mRNAs into categories based on previously obtained results (Figure 1C). **XUT/SUT** RNAs are typical PTC-containing NMD targets, as they have spurious short open reading frames followed by long untranslated regions (Malabat *et al*, 2015). The NMD targets group also comprise genuine **PTC**-containing protein coding mRNAs (He *et al*, 2003; Heyer & Moore, 2016) and mRNAs stabilized more than 2-fold upon Upf1 deletion (Figure S1A). Genes that express mRNAs without an obvious PTC were annotated **noPTC**, provided they express less than 10% of NMD-sensitive 5’ transcript isoforms and are not stabilized upon Upf1 deletion. Genes expressing more than 10% of NMD-sensitive 5’ transcript isoforms were annotated as possible PTC (**PTC_pos**). The remaining transcripts were considered to have an “**unknown**” NMD status. Genes corresponding to each category are individually listed in Table S1.

The RIPseq data were classified according the categories defined in Figure 1C. 2452 detected transcripts corresponded to **XUT/SUT**, and 465 to **PTC**. From the noPTC categories 345 were enriched with Upf1 (**noPTC_enr**, red), whereas 1762 were not (**noPTC** pink - Figure 1D). The **noPTC_enr** transcripts, were stabilized at the similar levels than other noPTC mRNA upon Upf1 deletion (Figure S1B). Examples of the PTC category were strongly enriched in Upf1 immunoprecipitates (Figure 1E). Among the noPTC representative examples, some were enriched, though to lower levels than the PTC ones. Noteworthy, highly expressed genes of the noPTC_enr category such as ribosomal protein-coding mRNA or histone-coding transcripts represent an important fraction of Upf1-associated mRNA (Figure 1B, 1E and Table S1).

mRNAs stability relies on their poly(A) and translation status (Lima *et al*, 2017). The codon-optimality was determined as the percentage of optimal codon in each coding sequence (Pechmann & Frydman, 2013). The noPTC categories whether enriched or not with Upf1, have a higher codon optimality (Figure S1C) and longer half-lives (Miller *et al*, 2011) (Figure S1D). Optimal codon content correlated with short poly(A) tail lengths determined by (Subtelny *et al*, 2014) (Figure S1E). In agreement with previous reports (Audebert *et al*, 2024), the most abundant transcripts showed the highest proportion of optimized codon and the shortest poly(A) tails.

We conclude that high enrichment of NMD targets in association with Upf1 is accompanied by enrichment of mRNAs that do not contain a PTC (noPTC) and for which no link with NMD has been previously established.

### Nanopore Direct RNA sequencing redefines Upf1-bound mRNAs landscape

We identified a stable and abundant population of transcripts that lack PTCs and are robustly associated with Upf1. Notably, their levels remained unchanged upon Upf1 deletion (Figure S1A and S1B). We hypothesized that the population of Upf1-associated transcripts might represent a specific subfraction of mRNAs for each of the hundreds of corresponding genes. Direct RNA sequencing (DRS) using Oxford Nanopore Technology enables the sequencing of individual mRNA molecules from their poly(A) tails to their 5′ ends (Workman *et al*, 2019), providing poly(A) tail length information for each transcript without introducing biases associated with PCR amplification (Figure 2A). We thus determined by DRS the transcripts enriched in the Upf1-purified fraction compared to the WT input and evaluated their poly(A) tail lengths. Unlike Illumina sequencing, DRS also distinguishes specific RNA isoforms.

**Figure 2.**
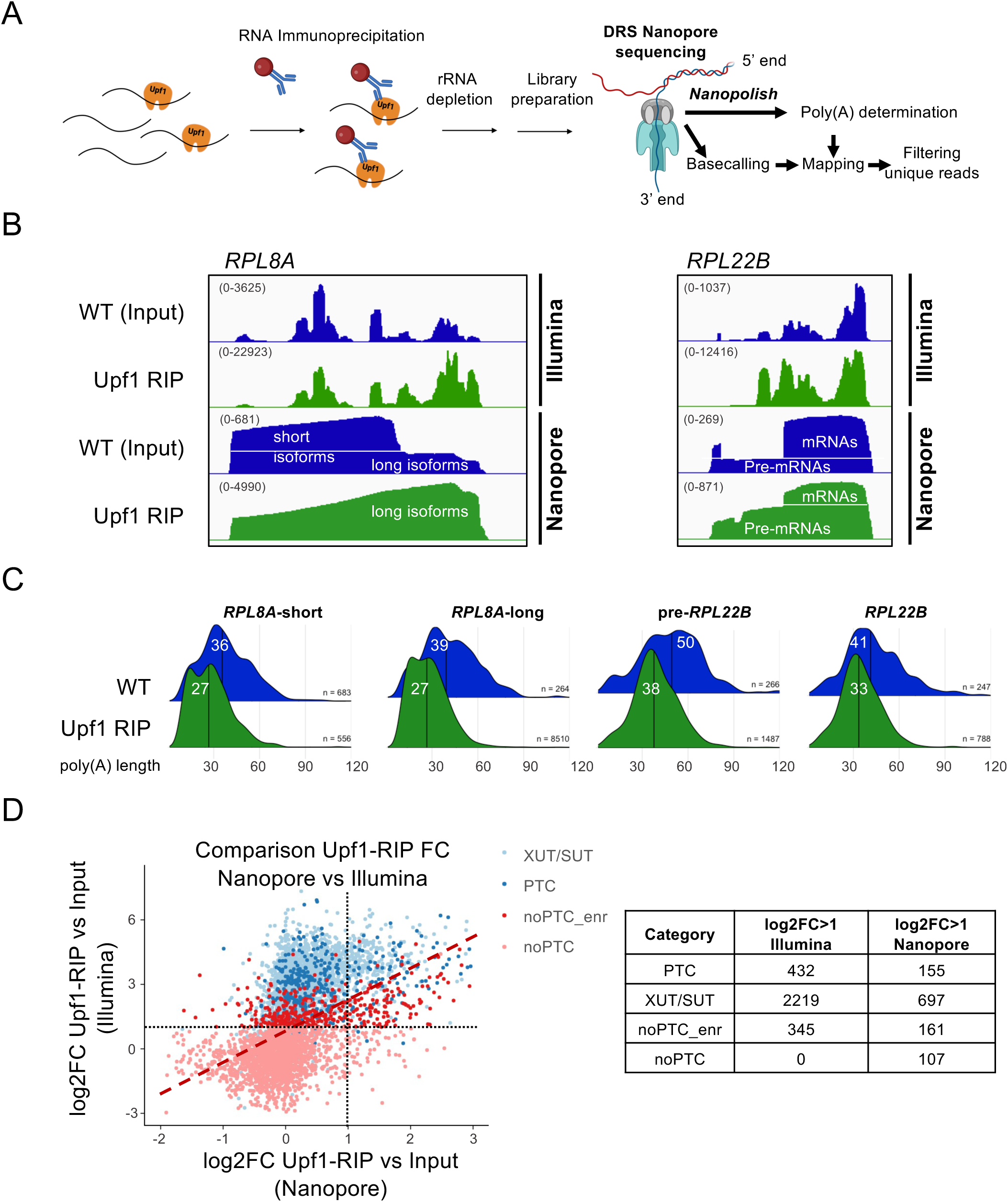
Nanopore Direct RNA sequencing redefines Upf1-bound mRNAs landscape. (A) Schematic representation of RNA-Immunoprecipitation (RIP) followed by Nanopore sequencing. (B) Signal intensity of RNA associated with Upf1-TAP (green) compared to total RNA from WT (blue) for Illumina or Nanopore sequencing (with Autoscale option in the Integrated Genome Viewer, IGV). Two examples are shown: *RPL8A*, which generates short and long 3’ UTR isoforms, and *RPL22B*, which can exist as pre-mRNA (pre-*RPL22B*) or mature mRNA (*RPL22B*). The scale is indicated in brackets. (C) Poly(A) tail lengths profiles of RNA associated with Upf1-TAP (green) compared to total RNA from WT (blue) for *RPL8A* and *RPL22B* isoforms. Poly(A) tails were determined using Nanopolish (Workman et al., 2019). The median poly(A) tail is indicated. (D) Scatterplot comparing log2FC between Upf1-RIP and WT in Nanopore (abscissa) and Illumina (ordinate), for transcripts from PTC: XUT/SUT and PTC, and noPTC categories: noPTC Upf1-enriched (noPTC-enr) and not enriched (noPTC). Dotted lines (black) indicate a 2-fold threshold (log2FC = 1). The dashed red line represents the linear regression trend line (Pearson’s correlation coefficient, r = 0.48072 p value < 2.2 10−16). The number of transcripts with a log2FC between Upf1-RIP and WT >1 are listed in a table (left panel). See also Tables S3 and S4.

For instance, *RPL8A* mRNA exists in both long and short isoforms that are difficult to distinguish based on Illumina sequencing, but are clearly distinguished with Nanopore. Interestingly, DRS revealed a stronger association of the long isoform of *RPL8A* with Upf1 than that of the short isoform, despite being a minority in the WT Input sample (264 “long” *vs* 683 “short” reads, Figure 2B, left panel). Quantitative analysis of the raw data showed a 32.2-fold relative enrichment of the *RPL8A-long* isoform in the Upf1-purified sample compared to the WT Input (8510 *vs* 264 reads). In contrast, the *RPL8A-short* isoform was proportionally not enriched in these conditions (0.81-fold, 556 *vs* 683 reads), suggesting the recognition of the long isoform by Upf1. This difference was not correlated with the median poly(A) tail lengths, similar for the two isoforms in the input and purified fractions (Figure 2C, left panels). Similarly, DRS enabled the quantification of the relative proportions of *pre-RPL22B* and mature *RPL22B* mRNAs. However, in the WT input sample, the poly(A) tail profiles of *RPL22B* and *pre*-*RPL22B* differed, with median poly(A) tail lengths of 41 and 50 As, respectively. In the Upf1-purified fraction, these lengths were further reduced to 33 and 38 As, respectively. Notably, the PTC-containing *pre-RPL22B* isoform exhibited a longer poly(A) tail overall compared to the mature *RPL22B* isoform lacking a PTC (Figure 2C, right panels).

To evaluate the overall differences between Illumina and Nanopore quantifications, we plotted log2 fold changes (log2FC) calculated from DESeq2-normalized counts in both datasets (Figure 2D). Despite the biases inherent in both techniques (PCR amplification for Illumina and oligo(dT) selection for Nanopore), the overall trend revealed a positive correlation, indicating a concordance between the two sequencing technologies. However, the number of transcripts classified as enriched (log2FC > 1) from Nanopore sequencing was notably reduced in the PTC categories (PTC and XUT/SUT) and the noPTC_Upf1-enriched category (noPTC_enr). Only the noPTC category, which by definition was not enriched in Upf1-RIP in Illumina sequencing, displayed more transcripts enriched in Nanopore sequencing (see Table Figure 2D). We conclude that Nanopore sequencing provide a distinct quantification of Upf1-bound RNAs while also defining isoform complexity. It allows the concomitant investigation of enrichment and poly(A) status in the Upf1-associated mRNAs, crucial for the next step of our study.

### Most Upf1-associated mRNAs lack a PTC and have a short poly(A) tail

Two families of Upf1-associated transcripts (PTC and noPTC_enr) emerged from short read Illumina sequencing, each characterized by distinct properties such as codon optimality and half-lives (Figure S1C-D). The global poly(A) tail distribution of immunoprecipitated RNAs obtained from Nanopore DRS using either Upf1 (Upf1-RIP) or eIF4G2 (eIF4G-RIP) were compared to distributions in a wild-type input (WT) or an *upf1*Δ RNA pool (Figure 3A). A tagged version of eIF4G2 - a subunit of the cytoplasmic cap-binding complex eIF4G (Goyer *et al*, 1993), which binds capped mRNAs (Schwartz & Parker, 2000), was used as a qualitative control for purification. The global poly(A) tail length distribution in the WT strain reflects the steady-state status of mRNA, encompassing both intact mRNA molecules and RNA fragments generated from the 5’ end, as sequencing is anchored at the poly(A) tail. Interestingly, transcripts in the WT sample exhibited a tri-modal distribution, with peaks at approximately 10 As, 30 As, and 55 As. In contrast, transcripts in the Upf1-RIP sample displayed only two major peaks (10 As and 30 As), while the eIF4G-RIP control primarily contained transcripts with longer poly(A) tails, showing a predominant peak at ∼55 As (Figure 3A). Since sequencing of eIF4G-bound mRNAs captured intact (capped) transcripts, and filtered mRNAs with extended poly(A) tails in all categories (Figure 3 and S2), it suggests that mRNA with shorter poly(A) might have in part lost their cap and could be degradation intermediates as they are depleted in eIF4G-RIP.

**Figure 3.**
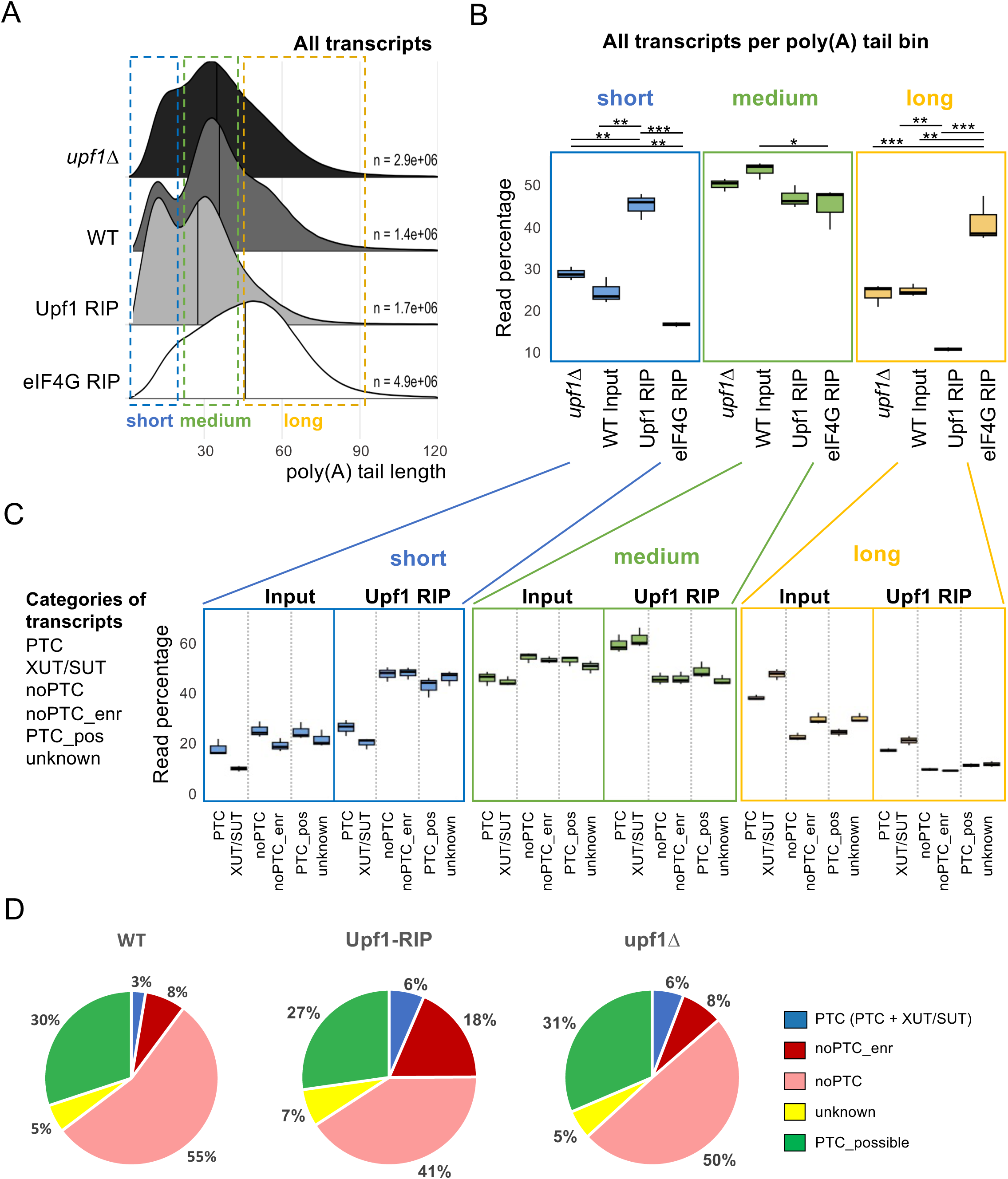
Most Upf1-associated mRNAs lack a PTC and have a short poly(A) tail. A) Ridge plot showing the global poly(A) tail length distribution across all transcripts from WT (Input) and *upf1*Δ total RNA, as well as Upf1-RIP and eIF4G2-RIP samples. The colored dotted sections indicate: short poly(A) tail (<25A, blue), medium poly(A) tail (>25A, <50A, green), long poly(A) tail (>50A, yellow). (B and C) Boxplots comparing the proportion of poly(A) tail length in short, medium and long bins across WT (Input) and Upf1-RIP samples transcriptome-wide (B) or comparing subcategories (PTC, SUT/XUT, noPTC, noPTC_enr, PTC_poss, and unknown) between Upf1-RIP and WT input RNAs (C). Pvalues (Tukey HSD test): ns = p> 0.05, * = p<0,05, ** = p<0,01, *** = p<0,001 and **** = p<0,0001. See also Table S2 for detailed pvalues. (D) Pie chart representing the proportion of each transcript category across the indicated samples: PTC containing NMD targets (PTC and XUT_SUT, blue); noPTC-Upf1enr (red), noPTC, (pink); unknown (yellow) and possible NMD (green). See also Table S1 and Figure S2.

To facilitate visualization and interpretation, reads were classified into three poly(A) tail length bins: short (0–25 A), medium (25–50 A), and long (>51 A) (Figure 3A and 3B). Strikingly, the proportion of transcripts with short poly(A) tails was significantly higher in the Upf1-RIP sample (44%) compared to the WT input (24%) (Figure 3B). This enrichment of short poly(A)-tailed transcripts was manifest regardless of the level of mRNA enrichment in the Upf1-RIP (noPTC_enr and noPTC categories, Figure 3C), mirroring the overall profile observed Figure 3B. The proportion of short poly(A) tails was likely underestimated across all the samples. Specifically, we used the standard Nanopore DRS library preparation kit that relies on oligo(dT)10 priming, which inherently biases against the mRNAs with shortest poly(A) tails. Furthermore, this increase in short poly(A)-tailed transcripts in the purified fraction was not due to a technical artifact, as RNA obtained under similar conditions from eIF4G-RIP experiments displayed extended poly(A) tails. In this sample, medium and long poly(A) transcripts represented 44% and 40% of the total, respectively, confirming a good RNA integrity in these experiments (eIF4G RIP; Figure 3A and 3B).

Together, these results indicate that, contrary to expectation and in addition to PTC transcripts, Upf1 is predominantly associated with a short-poly(A)-tailed subpopulation of noPTC transcripts.

The enrichment of short poly(A)-tailed mRNAs within the Upf1-RIP sample was unexpected, given that NMD is considered to be deadenylation-independent (Muhlrad & Parker, 1994, 1999; Cao & Parker, 2003). Furthermore, we recently demonstrated that NMD targets are inherently unstable and undergo a rapid degradation, which results in longer poly(A) tails (Audebert *et al*, 2024). Consistent with this prediction, we observed that PTC-containing NMD targets exhibited longer poly(A) tails compared to other transcript categories, both in the WT input (long tails) and in the Upf1-RIP sample (long and medium tails, Figure 3C). Representative PTC-containing NMD targets such as XUT/SUT or *RPL22B* pre-mRNA corroborated this trend (Figure S2A). However, Upf1-associated PTC-NMD targets exhibited overall slightly shorter poly(A) tails compared to the WT input (Figure S2B). While the targeting of NMD transcripts occurs independently of deadenylation, this observation suggests that a moderate deadenylation may take place before the resolution of NMD and Upf1 dissociation. In *upf1*Δ cells, NMD targets (XUT/SUT and PTC) displayed shortened poly(A) tails, similar to other categories of mRNA, and consistent with their stabilization in the absence of functional NMD (Figure S2C *upf1Δ vs* WT).

Given that DRS Nanopore technology directly sequences mRNA, with each read corresponding to a unique initial RNA molecule, multiple assignments were filtered out to keep the most relevant annotation for each read (see STAR method for details). We estimated the relative abundance of each transcript within different categories (e.g., PTC, noPTC etc.). Short poly(A) tails were significantly enriched across the noPTC noPTC-enr, PTC_pos and unknown categories, collectively accounting for 94% of the total reads in Upf1-RIP sample (Figure 3D), closely mirroring the overall distribution observed in Figure 3B. As anticipated, PTC-containing NMD targets constituted a small fraction in the WT input (3%), consistent with their rapid degradation by the NMD machinery. Interestingly, they accounted for only 6% in the Upf1-RIP and in the *upf1*Δ samples (+3%; Figure 3D). Importantly, we demonstrated that the presence of noPTC mRNAs in the Upf1-RIP fraction was not merely due to background noise, as the selected mRNA population during immunoprecipitation was clearly distinct from that in the input, when taking into account the poly(A) tail size distribution (Figure 3A-B and noPTC representative *HHT2* and *TDH3*, Figure S2A). These findings suggest that noPTC short poly(A)-tailed transcripts are a major and specific fraction of the mRNA bound by Upf1.

### Short poly(A)-tailed noPTC mRNAs accumulate in heavy polysome fractions

While the association of PTC-containing NMD substrates with Upf1 is correlated with their increased levels in the absence of NMD, noPTC transcripts are not known to be affected by such mutant conditions. This fact may be explained by the relatively low abundance of short poly(A) forms issued from noPTC mRNAs in comparison with the longer poly(A) forms. Thus, even if this fraction was affected by NMD, the impact of its stabilization in an NMD mutant on the global levels of the corresponding population of transcripts might be below usual detection thresholds. To test this hypothesis, and because NMD is a translation-dependent pathway, we first investigated the translation status of short-tailed mRNA. To do so, we fractionated mRNPs based on their sedimentation through sucrose gradients and analyzed the poly(A) tail lengths. We chose the *HHT2* transcript, a representative mRNA from the noPTC-Upf1-enriched (noPTC-enr) category that encodes one of the two copies of the core histone H3. Northern blotting experiments with probes specific of *HHT2* revealed that most *HHT2* mRNAs were localized in polysome fractions 1 to 3 in WT cells, despite the higher levels of 18S and 25S rRNAs in fractions 4 and 5. In *upf1*Δ cells, these mRNAs showed a more even distribution, from 1 to 5 (Figure 4A). To assess the poly(A) tail length of *HHT2* across polysome fractions, we employed the extended Poly(A) Test (ePAT), a method allowing to visualize the diversity of poly(A) tail lengths of specific mRNAs (Jänicke *et al*, 2012 and Figure 4B). Consistent with our Nanopore DRS results, no significant difference in global poly(A) tail length was detected between WT and *upf1*Δ RNA samples (Figure 4B-C, left panels). Interestingly, in both WT and *upf1*Δ conditions, *HHT2* transcripts with short poly(A) tails were enriched in the heavier polysome fractions (fractions 3 to 6 Figure 4A), suggesting an accumulation of short poly(A) forms of *HHT2* in these fractions. Remarkably, in *upf1*Δ, short poly(A) forms were also detected in the lighter polysome fractions (fractions 1 and 2 - Figure 4B-C). This differs from PTC-containing NMD targets that were found principally in the monosome fraction (Heyer & Moore, 2016). Finally, sucrose gradient fractionation followed by ePAT identified an Upf1-dependent shift in the size of mRNPs containing short poly(A)-tailed mRNAs, suggesting the involvement of NMD in their remodeling or translation.

**Figure 4.**
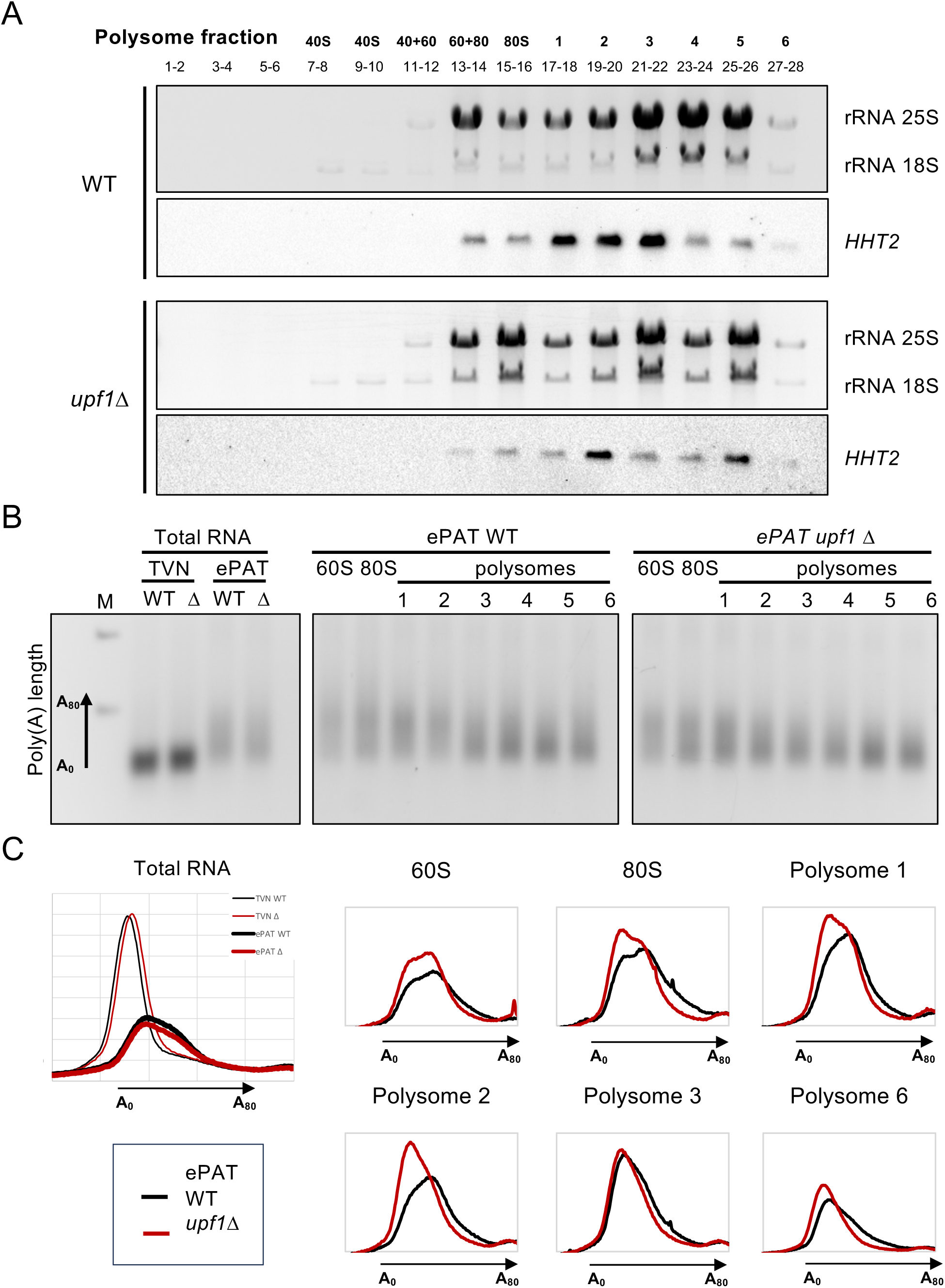
Short tailed noPTC mRNAs are present in heavy polysome fractions. A) Total RNA extracted from polysome fractions across sucrose gradient in wild-type or *upf1*Δ strains were visualized by ethidium bromide staining. *HHT2* signal was revealed by Northern blot (see also STAR methods). B) Extended Poly(A) Test (ePAT,^37^) and TVN-PAT (=A0) experiments on *HHT2* transcript issued of RNA extracted from sucrose gradient. Ethidium-stained agarose gel with the *HHT2* ePAT signal from Total RNA (left) or from the polysome fractions (right - corresponding to fractions Figure 4A) from WT and *upf1*Δ strains, flanked by a molecular weight marker (M). C) ImageJ integration of Total RNA ePAT signals (no fractionation – left panel) from wild-type (black) and upf1Δ (red) strains or from polysome fractions (as in Figure 4A – right). The abscissa shows the relative poly(A) tail length (nt), and the ordinate represents the integrated signal (arbitrary unit from ImageJ). See also Figure S3.

### Upf1 targets short poly(A)-tailed mRNAs for degradation

To separate RNAs based on their poly(A) tail length, we adapted a poly(A) fractionation protocol(Meijer *et al*, 2007) using oligo dT magnetic beads (NEB). This approach enables the recovery of RNAs without or with a short (S), medium (M) and long (L) poly(A) tails (Figure 5A and see STAR method). We confirmed the efficient separation of mRNA species with different poly(A) tail lengths across the three fractions using ePAT on *HHT2* (Figure 5B, WT). A significant portion of the signal was detected in the medium fraction, consistent with its relatively good stability (half-life of 15.3 minutes according to (Miller *et al*, 2011). We also tested the *RPL28* pre-mRNA, a well-known PTC-NMD substrate, in which most of the signal was found in the L fraction. This result aligns with our Nanopore sequencing findings and with the expected pattern of a predominantly long poly(A) tail for an unstable transcript (Figure 5B). In the *upf1*Δ samples, the *HHT2* ePAT signal in the S fraction was significantly increased compared to WT, while the other fractions, including total RNA, remained unaffected. This suggests that the deadenylated forms of *HHT2* are stabilized in the absence of Upf1. In contrast, the *RPL28* pre-mRNA NMD control showed a strong increase of the M fraction in *upf1*Δ, reflecting the expected stabilization of this NMD target, which leaves more time for deadenylation (Figure 5B).

**Figure 5.**
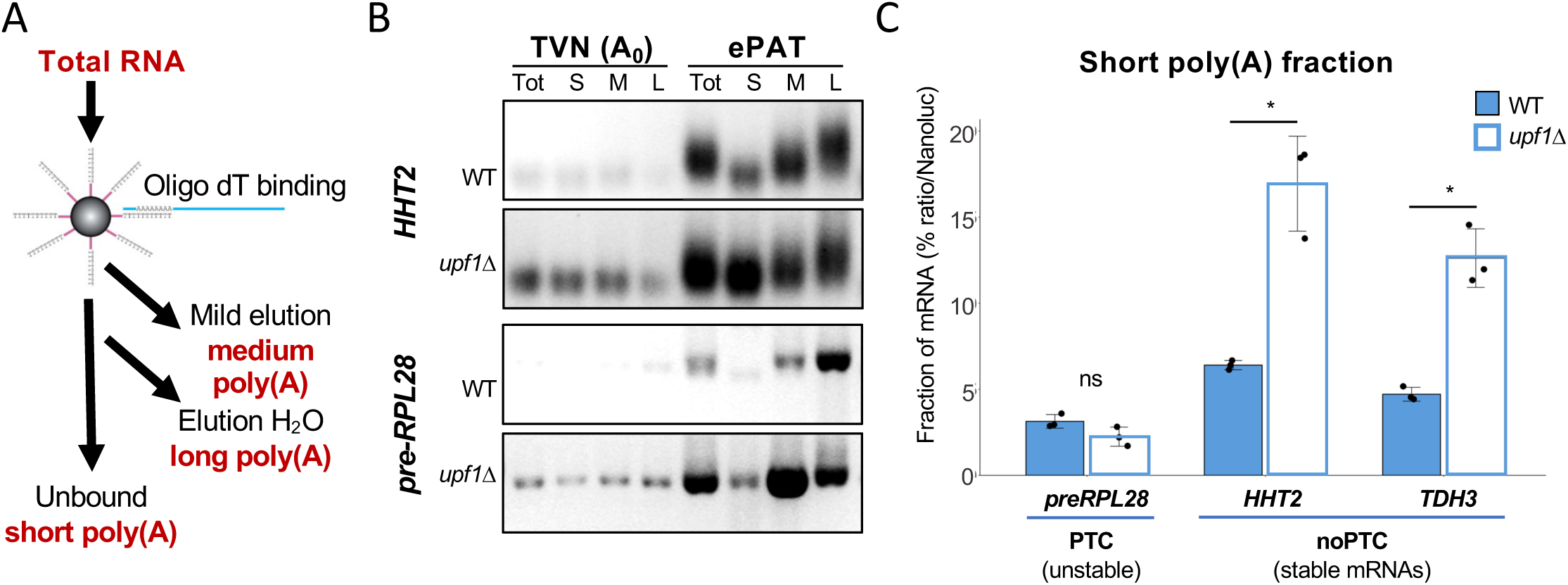
Upf1 targets short poly(A)-tailed mRNAs for degradation. (A) Experimental workflow for poly(A) tail length fractionation using oligodT beads (see also STAR methods). (B) Ethidium bromide-stained agarose gel showing ePAT and TVN-PAT (A0) results for *HHT2* and pre-*RPL28* across Total RNA (Tot), short (S), medium (M), and long (L) fractions in wild-type and *upf1*Δ strains. (C) Barplot showing the relative proportion of transcripts in the short poly(A) tail fraction for *preRPL28* (NMD control), *HHT2* (noPTC-Upf1enr) and *TDH3* (noPTC) transcripts, in wild-type (solid-bar) and *upf1*Δ strains (outlined-bar). Data represent the mean of three biological replicates; black dots indicate individual replicate values, and error bars represent the standard deviation (SD). Asterix* indicates significant p-values (t-test): *HHT2* p = 0.02124; TDH3 p = 0.01165). See also Figure S4.

Since ePAT experiments are not fully quantitative, we measured transcript levels in the fractions using RT-qPCR. To evaluate fractionation efficiency, we introduced a calibrated pool of *in vitro* transcribed Nanoluciferase mRNAs with varying poly(A) tail length (A0, A12, A30 and A50) prior to fractionation and estimated their distribution across the three poly(A) fractions by RT-qPCR (Figure S3A). As expected, A0 Nanoluciferase RNA was mostly found in the unbound (S) fraction, while A12 was mainly in the M, and A30 or A50 in the L fractions (Figure S3A).

We compared the relative amounts of representative transcripts in the different fractions: noPTC *HHT2* (enriched in Upf1-RIP) and *TDH3* (not enriched), PTC *RPL28* pre mRNA, and the *ScR1* ncRNA (used as a deadenylated control), in both WT and *upf1*Δ strains. The *ScR1* non-coding control was mainly found in the S fractions, while the PTC-NMD control *RPL28* premRNA was predominantly in the L fraction in the WT strain shifting to the M fraction in the *upf1*Δ strain, corroborating the ePAT results (Figure S3B). Interestingly, in the WT condition, for both *HHT2* and *TDH3* noPTC transcripts, the main signal was detected in the M fraction, suggesting that these transcripts naturally possess medium poly(A) tails at steady state. Moreover, a significant increase in the S fraction was measured for these mRNAs in the absence of Upf1 (Figure S3 and Figure 4C).

Taken together, these results support a role of Upf1 in the clearance of the short poly(A) tailed fraction of noPTC transcripts, regardless of their relative enrichment in the Upf1 purification.

### NMD stimulates the degradation of short poly(A) tailed mRNAs

To further characterize the role of Upf1 on short poly(A)-tailed mRNAs, we used the Tet-repressible *HIS3* codon optimized gene (*HIS3-opt*, Figure 6A) as a representative noPTC reporter in a reconstituted system. *HIS3-opt* gene is fully codon-optimized for translation and its mRNA is relatively stable(Audebert *et al*, 2024). The impact of Upf1 on *HIS3-opt* mRNA stability was evaluated by comparing changes in mRNA levels after transcriptional shutoff with doxycycline in WT and *upf1*Δ strains. Additionally, poly(A) RNA fractionation was performed to examine the distribution of *HIS3-opt* mRNA across total RNA samples and in the three sub-fractions (short, medium and long poly(A) tail - Figure 6A and STAR method). While no significant differences in mRNA half-lives were observed between WT and *upf1*Δ strains in the total RNA samples or in the medium and long sub-fractions, the half-life of *HIS3-opt* mRNA in the short fraction increased in the absence of Upf1 (24.4 minutes vs. 12.5 minutes in the WT). These findings confirmed the critical role of Upf1 in selectively targeting the short poly(A)-tailed sub-fraction of the *HIS3-opt* reporter for degradation.

**Figure 6.**
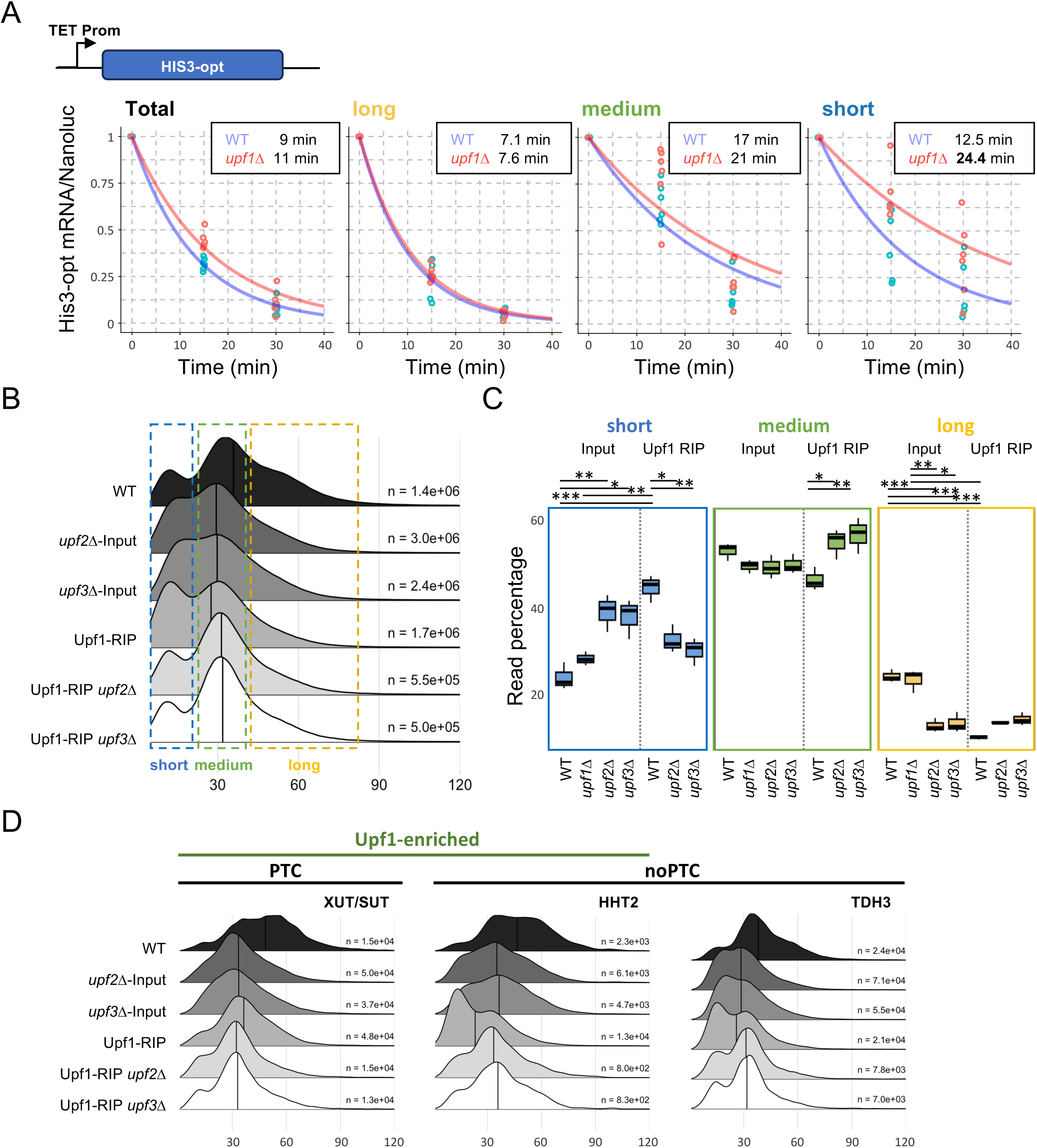
NMD stimulates the degradation of short poly(A)-tailed mRNAs. (A) Schematic representation of the HIS3-opt reporter. Kinetic of degradation of HIS3-opt reporter in wild-type (WT, blue) and upf1Δ (red) strains. Following a poly(A) fractionation as in Figure 4, the HIS3-opt reporter was quantified by RT-qPCR on RNA extracted from cells collected at t=0, t=15 and t=30 minutes after transcription shutoff. Normalization was performed using Nanoluciferase RNA as an internal control (see STAR methods). The half-life of HIS3-opt was calculated by fitting an exponential decay model to the data using a nonlinear least squares approach. The degradation constant (k) was estimated, and half-lives were computed as t_1/2_= ln(2)/k, are reported in boxes. Fitted exponential decay curves based on these calculations are displayed, and half-life values are reported in boxes. (B) Ridge plot showing the global poly(A) tail length distribution obtained by Nanopore sequencing across WT or *upf2*Δ and *upf3*Δ context, in input or Upf1 RIP samples. (C) Boxplot comparing the proportions of poly(A) tail length bins (short/medium/long) between Upf1-RIP and Input RNAs in WT, *upf2*Δ, and *upf3*Δ strains, alongside results from upf1Δ (as shown in Figure 2B). (D) Ridge plot depicting the poly(A) tail length distribution for transcripts in different categories: XUT/SUT (ncRNA), PTC-containing mRNAs, and specific examples such as *HHT2* (noPTC-Upf1enr) and TDH3 (noPTC). Distributions are shown for Upf1-RIP and Input RNAs in WT, upf2Δ and upf3Δ strains. See also Figure S4.

Compared to the global kinetic of degradation measured as the half-life in total RNA sample, the apparent half-lives of *HIS3-opt* in each sub-fraction results from two overlapping processes: deadenylation and degradation. Long-tailed *HIS3-opt* mRNAs have a shorter half-life than medium-tailed mRNAs. This difference is not due to a faster degradation but likely reflects rapid deadenylation, which transitions these mRNAs into the medium-tailed fraction. Similarly, stable medium-tailed mRNAs undergo further deadenylation and eventually shift into the short-tailed fraction. For the short poly(A) fraction, deadenylation no longer contributes to mRNA turnover as these transcripts cannot be further deadenylated. Therefore, the half-life measured in this fraction primarily reflects the effects of mRNA decay. These findings suggest that the stability of short poly(A)-tailed mRNAs critically depends on the presence of Upf1 in this reconstituted system (Figure 6A and see also Discussion).

Given that NMD can potentially target mRNAs as a result of individual ribosome frameshifting(He *et al*, 2003), we investigated the affinity of Upf1 for modified versions of the *HIS3-opt* reporter to address this possibility. These constructs included versions where possible stop codons in frame +1 and +2 were either removed (no stop) or retained a possible premature termination codon in frame +1 and in frame +2 (Stop +1+2 – see Figure S4A). To assess Upf1’s association with *HIS3-opt* mRNA we performed RT-qPCR on Upf1-RIP samples and WT input, normalizing the results to *TDH3*, a non-enriched but abundant endogenous mRNA (Figure S4B). The *HIS3-opt* mRNA was enriched 2.31 fold in Upf1-RIP compared to input, which is in the same range as the enrichment observed for *HHT2* (5.7 fold; Figure 1E). This enrichment indicates that Upf1 is associated with the *HIS3-opt* transcript in a comparable manner to that of other noPTC_enr mRNAs. We evaluated the affinity of the “no stop” reporter (with no possibility to find a PTC in a frameshifted phase), and the Stop +1+2 reporter (with a possibility to frameshift and find one PTC into other phases). The analysis of *HIS3-opt* reporter mRNA levels in the RIP fraction relative to total RNA input revealed similar enrichment across the *HIS3-opt*, no stop, and Stop +1+2 constructs (2.31, 2.75, and 2.72, respectively; Figure S4B). Therefore, the contribution of a frameshift effect for the tested transcript is negligible.

Having established that Upf1 is involved in the clearance of the short poly(A)-tailed population of noPTC mRNAs, we next investigated whether this effect depends on other NMD factors, since Upf1-mediated decay pathways independent of NMD factor have been described (Kim & Maquat, 2019). To assess the impact of NMD factors on Upf1’s role in the clearance of short poly(A)-tailed noPTC mRNAs, we conducted Nanopore DRS on Upf1-RIP fractions in *upf2*Δ or *upf3*Δ strains, and compared the results to their corresponding total RNA inputs (Figure 6B-D). As observed in the *upf1*Δ strain (Figure 3), the poly(A) profiles of inputs were largely comparable across WT, *upf2*Δ, and *upf3*Δ strains, but with a notable increase in short poly(A)-tailed mRNAs in *upf2*Δ and *upf3*Δ inputs. Strikingly, however, in the Upf1-RIP samples, the short fraction corresponding to short poly(A)-tailed mRNAs was drastically reduced in the *upf2*Δ and *upf3*Δ compared to the WT context (Figures 6B–D). These results demonstrate that Upf2 and Upf3 are required for the recognition and targeting of short poly(A)-tailed noPTC transcripts by Upf1.

Taken together, these findings highlight the critical role of Upf1 and the NMD machinery in the degradation of short poly(A)-tailed noPTC mRNAs. The use of a reporter system further validated Upf1’s functional role in clearing the short poly(A)-tailed fraction of stable noPTC mRNAs. Furthermore, our results demonstrated that Upf1 relies on the presence of key NMD cofactors, Upf2 and Upf3, to efficiently target the short poly(A)- tailed sub-fraction, strongly suggesting that short poly(A)-tailed degradation is mediated through a mechanism involving the full NMD machinery.

## DISCUSSION

Four decades of studies on Upf1 have highlighted its crucial role in shaping the transcriptome, primarily by targeting mRNAs with premature termination codons through NMD (Peltz *et al*, 1993; Muhlrad & Parker, 1999; Eberle *et al*, 2008). While the precise mechanism by which Upf1 detects these mRNAs remains an open question, PTC-containing reporters showed that Upf1 triggers mRNA degradation independent of deadenylation (Muhlrad & Parker, 1994, 1999; Cao & Parker, 2003), although it was not established if the same applies to all PTC transcripts. In this study, we comprehensively characterized Upf1 targets in *Saccharomyces cerevisiae* and found evidence that the NMD machinery also specifically targets short poly(A)-tailed mRNAs to facilitates their degradation, revealing a repertoire of NMD-regulated transcripts importantly larger than previously recognized.

First, we confirmed that Upf1 triggers PTC-containing mRNA degradation independently of deadenylation by observing longer poly(A) tails for PTC mRNAs in the Upf1-associated fraction (Figure 3). PTC-containing mRNAs are homogeneously recognized by the NMD. In contrast, short poly(A)-tailed mRNAs are a new population of transcripts that are specifically bound by Upf1 and was not detected until now, as they represent only a minor fraction within a heterogeneous population of "non-NMD mRNAs" (Heyer & Moore, 2016).

### Short poly(A) Upf1-targeted mRNAs are masked by mRNA heterogeneity

Nanopore sequencing revealed the specific affinity of Upf1 for short poly(A)-tailed transcripts. This discovery implies that enrichment values for a majority of transcripts in the Upf1 purified fraction based on Illumina sequencing are not truly representative, as they compare short poly(A)-tailed mRNAs with all poly(A) length isoforms for the mRNAs originating from a given gene. Despite the relative abundance of short poly(A)-tailed mRNAs in association with Upf1, largely surpassing the amounts of PTC-containing mRNA (Figure 3), these RNAs have not been considered until now as targets of NMD. To test the effect of inactivating NMD on short poly(A)-tailed mRNA stability, we fractionated cellular mRNA on the basis of the length of the poly(A) tails, which allowed us to identify a specific effect of Upf1 on short poly(A)-tailed mRNA levels and translation (Figure 4 and 5). Using a reconstituted noPTC *HIS3-opt* reporter, we identified variations in the intrinsic stabilities of short-, medium- and long-poly(A) tailed mRNA subpopulations, which would otherwise be indistinguishable in total RNA samples. Furthermore, we demonstrated that Upf1 and the NMD pathway significantly affect the levels (endogenous *HHT2* and *TDH3)* and stability (*HIS3-opt*) of noPTC short poly(A)-tailed mRNAs (Figures 5 and 6).

### A model for the clearance of short poly(A)-tailed mRNAs by NMD

Our results identified a subpopulation of transcripts that is characterized by short poly(A) tails and specific binding to Upf1. Moreover, NMD accelerates their degradation. These new findings lead to the obvious question about how Upf1 is recruited to short poly(A)-tailed mRNA. One of the simplest hypotheses is that a shortened poly(A) tail is accompanied by a decrease in Pab1 binding to the 3’ end of the transcript. By analogy with the “Faux 3’ UTR” NMD model, a lack of Pab1 in proximity of the stop codon would lead to a translation termination defect that provokes the recruitment of NMD factors, in particular Upf1. This hypothesis is consistent with our findings and previously published results (Eberle *et al*, 2008; Amrani *et al*, 2004).

Considering that a single Pab1 molecule binds approximately 25–30 adenines(Webster *et al*, 2018), we estimated the number of Pab1 molecules associated with each bins of poly(A) tail length in Nanopore data: short tails (<25 adenines) likely lack Pab1, medium tails (25–54 adenines) are bound by one Pab1 molecule, and long tails (>55 adenines) are associated with two or more Pab1 molecules. This suggests that short poly(A) tailed mRNAs, which are significantly enriched in the Upf1-associated fraction, may lack bound Pab1.

Although both targeted by the NMD, PTC and short poly(A) tailed noPTC transcripts have distinct behaviors. PTC transcripts are homogeneous, unstable and are likely charged with one or two Pab1 molecules (mainly medium and long tailed molecules in the WT Input – Figure 3 and S2). In addition, they have previously been shown to be enriched in the monosome fraction (Heyer & Moore, 2016), reflecting the short coding sequence and poor translation that are specific to these transcripts. In contrast, noPTC mRNAs are heterogeneous, stable, and most noPTC poly(A) forms are well translated, with only the short poly(A) tailed subpopulation that is targeted by the NMD. Short poly(A) tailed mRNAs are localized in heavy polysome, although if this localization corresponds to high levels of translation or to stalled ribosomes remains unclear. Short poly(A) tails accumulated in lighter polysome fractions in the absence of Upf1, consistent with a role of Upf1 in solving inefficient translation termination of short poly(A) tailed mRNAs (Figure 4). This profile could indicate either a high levels of translation or ribosome stalling at the stop codon, both scenarios not being mutually exclusive.

PTC and noPTC short poly(A) tailed NMD-targeted mRNAs are thus distinct. According to the “Faux 3’ UTR” model (*i.e.* PTC-NMD), the long distance between Pab1 and the stop codon dictates mRNA recognition by the NMD machinery, whereas in short poly(A)-NMD, it is the absence of Pab1. In both scenarios, translation termination is impaired due to the absence of Pab1’s stimulating effect on the translation termination factor eRF3 (Cosson *et al*, 2002). In both cases, Upf1 could be required to resolve the same translation termination defect. Altogether, our findings converge to a unified model in which short poly(A) tailed mRNAs are degraded via a specialized NMD-mediated pathway, which we term the "short poly(A) NMD pathway" (Figure 7).

**Figure 7.**
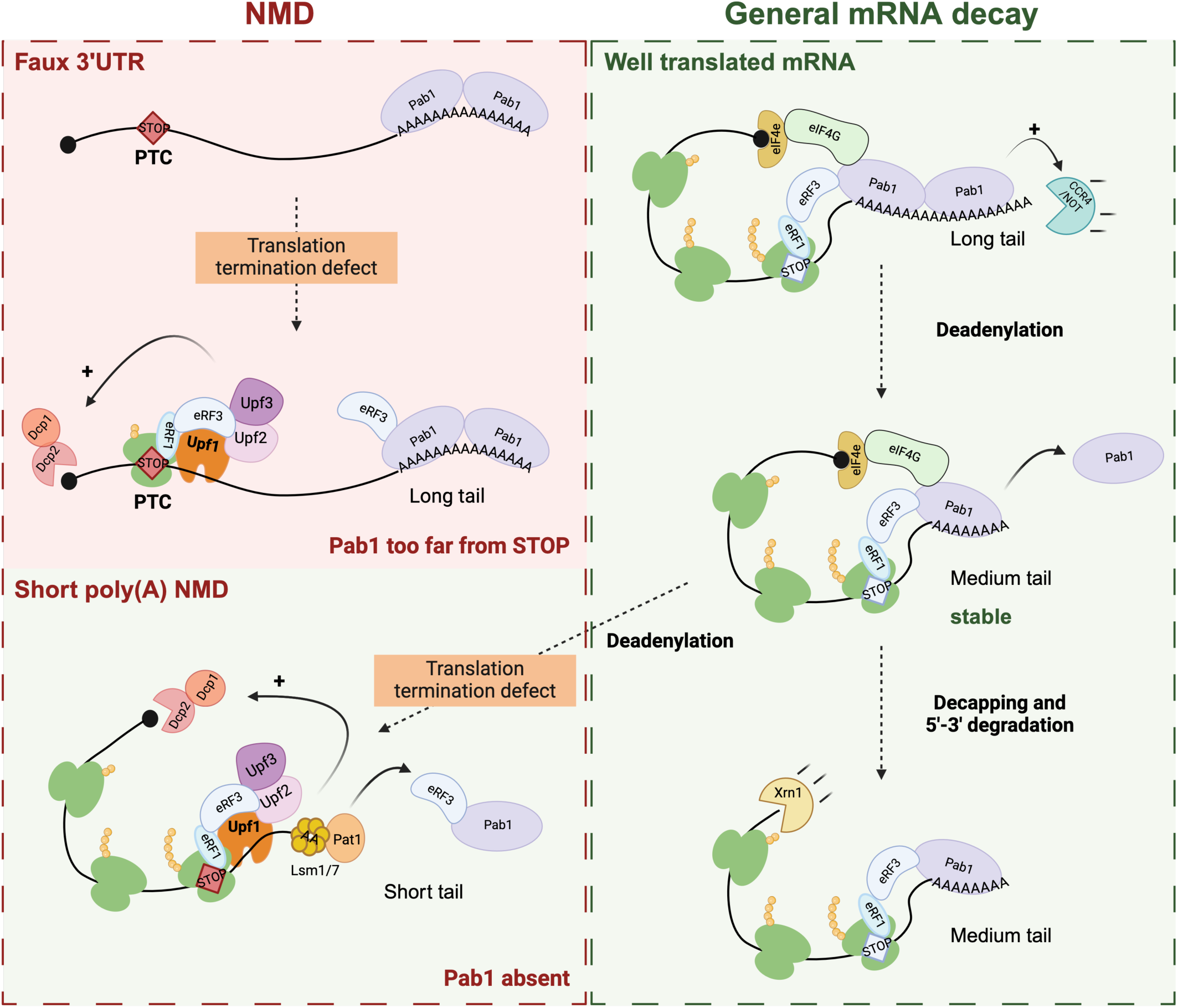
A model for the clearance of short poly(A)-tailed mRNAs by NMD. Schematic representation of classical **PTC-targeted NMD** (Faux 3’UTR model) and **short poly(A) NMD**, and its links with general mRNA decay.

Further validation of this model comes from the observation that Upf1-associated short poly(A) tailed mRNAs differ from Pab1-associated mRNAs. Specifically, we demonstrated that Upf1-associated noPTC mRNAs, including *HHT2*, exhibit short poly(A) tails. In the absence of Upf1, the short-tailed fraction of these transcripts is stabilized. Conversely, Pab1-associated *HHT2* transcripts present extended poly(A) tails (Audebert *et al*, 2024) suggesting that these two proteins target distinct subpopulations of the same transcript.

How could the short poly(A) NMD pathway be integrated into the current models of mRNA degradation following deadenylation? Oligoadenylated mRNAs are recognized by the Lsm1-7/Pat1 complex which stimulates decapping and subsequent mRNA degradation (Tharun *et al*, 2000; Bouveret *et al*, 2000; He & Parker, 2001). We previously identified the presence of both Upf1 and Lsm1-7/Pat1 complex in the same mRNPs (Dehecq *et al*, 2018). This initially seemed paradoxical as NMD is known to be deadenylation-independent and consequently occurs on mRNAs with long poly(A) tails (Muhlrad & Parker, 1994, 1999; Cao & Parker, 2003) (and also as shown in this study Figure 3). We conciliated this observation in light of our current findings: as Upf1 targets short poly(A) tailed noPTC mRNAs intended for general decay it is plausible that these transcripts can also be detected by the Lsm1-7/Pat1 complex.

The model described here in yeast is compatible with previous observations in other species. Alongside, the global poly(A) status of mRNA decay intermediates associated with the phosphorylated form of human Upf1 -the active form in humans - revealed a very short global poly(A) profile, possibly reflecting similar short poly(A)-tailed targets in humans (Kurosaki *et al*, 2018).

Finally, it would be valuable to determine the proportion of mRNA that are decapped and “normally” degraded before the last Pab1 is removed, in comparison to those targeted by short poly(A) NMD. This comparison would provide a more comprehensive estimate of the overall contribution of short poly(A) NMD to general mRNA decay and its broader impact on the transcriptome. Future research will be essential to deepen our understanding of the significance of poly(A) tail length as a regulatory feature in mRNA degradation, as well as to clarify the specific involvement of Upf1 in this aspect of mRNA regulation. This could conduct in understanding how NMD shapes the transcriptome in eukaryotic systems beyond its original characterization in the degradation of PTC-containing transcripts.

### Limitations of the study

Due to the priming step with an oligo-d(T)_10_ in standards Nanopore direct RNA sequencing protocols, fully deadenylated mRNA are counter selected. The very short poly(A) and deadenylated fractions are consequently probably underestimated. Considering the importance of the short poly(A)-tailed fraction of mRNA for certain transcripts in our results, it would be valuable to circumvent this bias in futures studies in order to depict a more accurate view of Upf1-target landscape. This could be done by using TERA3 for example (Ibrahim *et al*, 2021). In addition, the 5’ end status (cap or 5’P) could also not be determined by standard Nanopore sequencing, but it would be interesting to evaluate the fraction of Upf1 targets which are full length and capped, vs degradation products.

## ACKNOWLEDGMENTS

We thank Hugo Varet and Dina Kari for bioinformatic guidance. We thank Maïté Gomard, Alexandre Cornille, Estelle Gauthier and other GIM members for technical help, Lucia Oreus for media preparation and Hervé Le Hir, Olivier Bensaude and Micheline Fromont for helpful discussions and their critical review of the manuscript. This research was funded by the “Centre National de la recherche scientifique” and by the “Agence Nationale de la Recherche” (grants ANR-22-CE12-0004-01 - GB and ANR-18-CE11-0003 - CS). TG was also supported by ANR-17-CE11-0049-01 and the ANR-17-CE12-0024-02.

## AUTHOR CONTRIBUTIONS

Conceptualization TG, CS and GB; Methodology CS and GB; Software TG and GB; Formal Analysis TG and GB; Investigation TG, and GB; Resources CS and GB; Data Curation GB; Writing – Original Draft TG and GB; Writing –Review & Editing TG, CS and GB; Visualization TG and GB; Supervision and Project Administration GB; Funding Acquisition CS and GB.

## DECLARATION OF INTERESTS

The authors declare no competing interests.

## SUPPLEMENTAL INFORMATION

Supplemental Information includes 4 figures, and 4 tables and can be found online

Document S1. Figures S1 to S4

Table S1. List the yeast transcript features with coordinates and categories. Excel file containing additional data, related to Figure 1

Table S2. Excel file containing all the replicate values and pvalues, related to Figure 1, 3, 5, 6 and supplemental Figures S1 to S4.

TableS3, related to Figure 1, shows the normalized counts of Illumina sequencing data.

TableS4, related to Figure 3, shows the normalized counts of Nanopore sequencing data.

## STAR★METHODS

### RESOURCE AVAILABILITY

#### Lead contact

Further information and requests for resources and reagents should be directed to and will be fulfilled by the lead contact, Gwenael Badis.

#### Materials availability

Strain and plasmids generated in this study are available upon request from the lead contact

#### Data and code availability

Sequencing data reported in this paper has been deposited at GEO: GSE283053 (Illumina) and GSE284490 (Nanopore) and are publicly available as of the date of publication.

All original codes have been deposited at Zenodo at 10.5281/zenodo.14615891 and is publicly available as of the date of publication.

Any additional information required to reanalyze the data reported in this paper is available from the lead contact upon request.

### KEY RESOURCES TABLE

The items in the key resources table (KRT) must also be reported alongside the description of their use in the method details section. Literature cited within the KRT must be included in the references list. Please **do not edit the headings or add custom headings or subheadings** to the KRT. We highly recommend using RRIDs (see https://scicrunch.org/resources) as the identifier for antibodies and model organisms in the KRT. To create the KRT, please use the template below or the <u>KRT webform</u>. See the more detailed <u>Word table template</u> document for examples of how to list items.

| REAGENT or RESOURCE | SOURCE | IDENTIFIER |
| --- | --- | --- |
| **Antibodies** |  |  |
| Sheep anti-digoxigenin-POD (poly), Fab fragments | Roche | Cat#11633716001 |
| Mouse anti-FLAG M2 - HRP | Sigma-Aldrich | Cat#A8592 |
| Rabbit Peroxidase Anti-Peroxidase soluble complex | Sigma-Aldrich | Cat#P1291 |
| **Bacterial strains** |  |  |
| NEB 10-beta competent E. coli | New England Biolabs | Cat#C3019H |
| **Chemicals, peptides, and recombinant proteins** |  |  |
| Clarity Western ECL substrate | Bio-Rad | Cat#1705061 |
| SsoAdvanced Universal SYBR Green Supermix | Bio-Rad | Cat#1725271 |
| TruSeq Stranded mRNA sequencing kit | Illumina | Cat#20020594 |
| NextSeq500 flowcell | Illumina | Cat#20024906 |
| BamHI | New England Biolabs | Cat#R0136S |
| PstI | New England Biolabs | Cat#R0140S |
| Oligo d(T)25 magnetic beads | New England Biolabs | Cat# S1419S |
| Q5 high-fidelity DNA polymerase | New England Biolabs | Cat#M0491 |
| DNA polymerase I, Large (Klenow) Fragment | New England Biolabs | Cat#M0212S |
| T4 DNA ligase | New England Biolabs | Cat#M0202T |
| Direct RNA Sequencing Kit | Oxford Nanopore | Cat#SQK-RNA002 |
| RNasin ribonuclease inhibitor | Promega | Cat#N2511 |
| Complete protease inhibitor mix (no EDTA) | Roche | Cat#11873580001 |
| DIG RNA labelling kit (SP6/T7) | Roche | Cat#11175025910 |
| Digoxigenin-11-dUTP | Roche | Cat#11093088910 |
| Indole-3-acetic acid (IAA) | Sigma-Aldrich | Cat#I2886 |
| Beta-estradiol | Sigma-Aldrich | Cat#E2758 |
| Doxycycline | Sigma-Aldrich | Cat#D3447 |
| Rabbit IgG | Sigma-Aldrich | Cat#I5006 |
| DNase TURBO | ThermoFisher | Cat#AM2238 |
| Phusion high-fidelity DNA polymerase | ThermoFisher | Cat#F530L |
| Superscript II reverse transcriptase | ThermoFisher | Cat#18064022 |
| Superscript III reverse transcriptase | ThermoFisher | Cat#18080044 |
| Dynabeads M-270 epoxy | ThermoFisher | Cat#14302D |
| **Deposited data** |  |  |
| UPF1_RIP, WT and upf1Δ Truseq Illumina sequencing | GEO | GSE283053 |
| UPF1-RIP, WT input, upf1Δ eIF4G2-RIP, UPF1-RIP in upf2Δ (input and RIP) and UPF1-RIP in upf3Δ (input and RIP) Nanopore Direct RNA sequencing | GEO | GSE284490 |
| R scripts for filtering Nanopore DRS reads and generating R figures | Zenodo | 10.5281/zenodo.14615891 |
| **Experimental models: Organisms/strains** |  |  |
| LMA1667 - BY4741 *nam7(upf1)Δ::KANMX6* | (Giaever et al, 2002) | N/A |
| LMA2154 (BY4741) (*S. cerevisiae*) *MATa ura3Δ0 his3Δ1 leu2Δ0 met15Δ0* | (Brachmann et al, 1998) | N/A |
| LMA2194 - BY4741 + *NAM7(UPF1)::HIS3MX6* | (Ghaemmaghami et al, 2003) | N/A |
| LMA3730 – BY4741 + NAM7(*UPF1*)-HTP | This study | N/A |
| LMA3314 – BY4741 + TIF4632-TAP::HIS3MX6 | (Ghaemmaghami et al, 2003) | N/A |
| LMA3682 – PB1623 (*S. pombe*) + *nam7(upf1)*-CRAC | This study | N/A |
| **Oligonucleotides** |  |  |
| PCR primers | This study | Table_Primers |
| **Recombinant DNA** |  |  |
| pCM189-NFLAG-HIS3-100 p1640 (TETO7-NFLAG-HIS3-100) (URA3) | Audebert et al. 2024 | N/A |
| pCM189-NFLAG-HIS3-100-NoSTOP p1803 (TETO7-NFLAG-HIS3-100) (URA3) | This study | N/A |
| pCM189-NFLAG-HIS3-100-STOP+1+2 p1804 (TETO7-NFLAG-HIS3-100) (URA3) | This study | N/A |
| **Software and algorithms** |  |  |
| ImageJ (1.54) | National Institutes of Health (NIH)<br>https://imagej.net/ij/ | N/A |
| Image Lab Software (6.1) | Bio-Rad | 12012931 |
| CFX Maestro Software (qPCR) | Bio-Rad | 12013758 |
| STAR (2.7) | (Dobin et al, 2013) | N/A |
| Integrated Genome Viewer (IGV, 2.11) | (Robinson et al, 2011) | N/A |
| R (3.8-4.4) | (R Core Team, 2024)<br>https://www.R-project.org/ | N/A |
| RStudio | https://posit.co/products/open-source/rstudio/ | N/A |
| Guppy | Oxford Nanopore | N/A |
| Minimap2 (2.1) | (Li, 2018) | N/A |
| Samtools (1.9) | (Danecek et al, 2021) | N/A |
| Nanopolish (0.13.2) | (Workman et al, 2019) | N/A |
| Rscripts were deposited on Zenodo (see Deposited Data) | Zenodo | 10.5281/zenodo.14615891 |
| **Other** |  |  |
| ChemiDoc XRS+ imaging device | Bio-Rad | N/A |
| Trans-Blot Turbo transfer system | Bio-Rad | Cat#1704150 |
| Supported nitrocellulose membrane | Bio-Rad | Cat#1620094 |
| Amersham Hybond-N+ | Cytiva Life Sciences | Cat#RPN303B |
| NextSeq 500 | Illumina | N/A |
| Gel and PCR clean-up | Macherey-Nagel | Cat#740609.250 |
| MinION sequencing device | Oxford Nanopore | N/A |
| GridION sequencing device | Oxford Nanopore | N/A |
| NuPAGE 4–12% Bis-Tris electrophoresis gel | ThermoFisher | Cat#NP0323BOX |

### EXPERIMENTAL MODEL AND STUDY PARTICIPANT DETAILS

#### Yeast and Bacterial strain

All *Saccharomyces cerevisiæ* strains were derived from the BY4741 strain (leu2Δ0, his3Δ1, ura3Δ0, met15Δ0, MATa), obtained from the Euroscarf deletion collection (http://www.euroscarf.de) or from the TAP-fusion library (Ghaemmaghami *et al*, 2003). New yeast strains were generated by lithium acetate-based transformation of BY4741 strain with a PCR product containing a selection marker cassette flanked by 50 bp arms located upstream and downstream the targeted ORF.

Upf1(Ymr080w)-TAP strain LMA2194 (MATa, ura3Δ0, his3Δ1, leu2Δ0, met15Δ0, YMR080W-TAP:HIS3MX6) and eIF4G2(Ygl049c)-TAP strain LMA3314 (MATa, ura3Δ0, his3Δ1, leu2Δ0, met15Δ0, YGL049C-TAP:HIS3MX6) were obtained from the *S. cerevisiæ* TAP fusion library (Ghaemmaghami *et al*, 2003).

The Ymr080w-HTP strain LMA3730 used for the Illumina RIPseq experiment was obtained by transformation of the Ymr080w-TAP LMA2194 strain with the HpaI linearized plasmid pGIM1287 TAP2CRAC.

Escherichia coli strain NEB 10-beta (NEB Cat# C3019) was used for plasmid construction and amplification. All the plasmid inserts were verified by sequencing.

C-terminal TAP-tagged strains originated from the (Ghaemmaghami *et al*, 2003) collection. Deletions were tested by PCR amplification of the modified locus.

#### Growth conditions and media

##### Yeast

Yeast strains were grown at 30°C in YPD (20 g/L glucose, 10 g/L yeast extract, 20 g/L bacto-peptone, and 20 g/L bacto-agar for plates only) or in synthetic media without uracil to select transformants and maintain plasmids with the URA3 marker. Cultures were grown in exponential phase at OD_600_ of 0.6 – 0.8.

##### Bacteria

Escherichia coli strains were grown in LB medium at 30°C to prevent TetO7 promoter pop-out. The medium was supplemented with Ampicillin (50µg/ml) for plasmid selection.

### METHOD DETAILS

#### Criteria of gene category status assignment

mRNAs were first sorted based on their propensity to generate NMD-sensitive RNA isoforms with alternative 5’ends according to Malabat *et al*.(Malabat *et al*, 2015). mRNAs homogeneous in 5’ (**noPTC**) were defined arbitrarily as generating less than 10% of NMD-sensitive 5’ end isoforms, whereas those producing more than 10% of NMD-sensitive 5’ end isoforms were considered **possible-PTC**. In order to refine this assignment, mRNAs were reassigned **PTC** if they were either stabilized more than two-fold upon *upf1Δ (*visualized as a MAplot Figure S1A) or assigned direct NMD target (classes A and B in Heyer and Moore, 2016). The **unknown** category also corresponded to mRNA not present in other categories or for which we did not have enough information from our or previous datasets.

#### Plasmids construction

pCM189-HIS3opt plasmid (pCM189-NFLAG-HIS3-100 - TETO7-NFLAG-HIS3-100 #GIM1640), originally described by (Audebert *et al*, 2024) was modified by replacing the original HIS3-opt coding sequence by a Bam HI (NEB Cat# R0136S]) and PstI (NEB Cat# R0140S) digested fragment with all internal out-of-frame stop codons removed (No-Stop) or with a unique stop codon in frame +1 and +2 (Stop +1+2). The modified HIS3 sequences were cloned into pCM189 by ligation with T4 DNA ligase (NEB Cat#M0202T), amplified in Escherichia coli and sequenced (Eurofins genomic) generating plasmid pCM189-HIS3nostop (#GIM1803) and pCM189-HIS3stop12(#GIM1804).

#### RNA immunoprecipitation

TAP-tagged Upf1 containing yeasts were grown at 30°C in 2L YPD in exponential phase (OD_600_ of 0.6-0.8). Cells were centrifuged, rinsed with cold water and immediately frozen at −80°C until lysis. Yeast cells were lysed under FastPrep mixing in lysis buffer (20 mM Hepes-K pH 7.4, 100 mM KOAc, 10 mM MgCl2, RNasin 1/1000 (Promega Cat#N2511), Protease inhibitor (Roche Cat# 11836170001) using acid-washed glass beads (program yeast, *Saccharomyces cerevisiæ*, 6.0 m/sec / Quickprep / 40 sec 2 times). Lysates were centrifuged 10 min at 4000 rpm at 4°C, and the supernatant was collected. Upf1-TAP associated RNA were immunoprecipitated using Dynabeads M-270 epoxy (ThermoFisher Cat# 14302D) coated with Rabbit IgG (Sigma-Aldrich Cat# I5006) in lysis buffer (as described in Namane & Saveanu, 2022) supplemented with 20% Triton X-100. The lysate was incubated for 1.5 hours at 4°C with gentle agitation. After incubation, the supernatant was discarded, and stringent washing steps were performed to ensure specific binding using lysis buffer without protease inhibitors. Elution was carried out in TES buffer (2% SDS, 1X TE, 30 mM EDTA) at 65°C for 15 min. RNAs were extracted from the supernatant containing the Upf1-immunoprecipitated RNAs using phenol/chloroform and precipitated with ammonium acetate.

#### Reverse Transcription and quantitative PCR

RNAs were extracted from yeast after three phenol extraction (Total RNAs) or one (RIP). Contaminating genomic DNA was removed from 50 µg of total RNA by DNase treatment using 10 unit of Turbo DNase (Invitrogen Cat# AM2239) at 37°C for 20 min and DNAse was eliminated by incubating 5 min with the Inactivation Reagent following the manufacturer’s instructions. Priming was performed using random hexamers, denaturated at 65°C for 5 min before being placed in ice. Reverse transcription was carried out using SuperScript II Reverse Transcriptase (Invitrogen Cat# 18064022) in a mix containing 5X First-Strand Buffer, 0.1M DTT, and 10mM dNTPs. The reaction was incubated at 42°C for 45 min and 70°C 10 min for heat inactivation. Targeted transcripts were amplified using specific primers (at 5 µM final concentration see Primer Table for description) in a reaction mixture containing SYBR Green Mix (Bio-Rad Cat# 1725272) and incubated 95°C 10 min; 40 cycles (95°C 10 sec; 60°C for 30 sec).

#### RNA fractionation by poly(A) tail length

To fractionate RNA by poly(A) length, we adapted (Meijer *et al*, 2007) protocol, utilizing Oligo d(T)_25_ magnetic beads (NEB) and sequential elution steps. 50 to 100 µg of total RNA were incubated in 100 µL of 2X Binding Buffer (40 mM Tris-HCl pH 7.5, 1 M LiCl, 2 mM EDTA) at 65°C for 2 min, then 100 µL of prepared Oligo d(T)_25_ magnetic beads were added to the mixture and incubated for 15 min at RT, placed on a magnet, and the supernatant, containing unbound RNA (short or A-tails), was collected and precipitated using ammonium acetate. The beads were washed twice with Washing Buffer (20 mM Tris-HCl pH 7.5, 0.5 M LiCl, 1 mM EDTA); once with Low Salt Washing Buffer (20 mM Tris-HCl pH 7.5, 0.2 M LiCl, 1 mM EDTA). Two successive elution were performed with 1) 20 µL of 0.075X SSC buffer for 10 min at RT (medium polyA tails) and 2) 20 µL of nuclease-free water for 10 min at RT (long polyA tails).

#### Sucrose gradient for polysome fractionation

Sucrose gradient of wild-type and *upf1*Δ yeast extracts were performed as described in (Tesina *et al*, 2019), except that RNAs (not proteins) were extracted from each fraction by phenol/chloroform method, precipitated using ammonium acetate and resuspended in H2O. RNA fractionation was controlled loading half of each fraction on a 1% agarose gel to assign each fraction to a status (40S, 60S, 80S and polysomes 1 to 6 and Figure S3).

#### Northern Blot

Northern blot experiments were performed using a PCR with GB1679 and GB1690 primers and DIG RNA Labelling kit (SP6/T7), Roche (Cat#11175025910) for RNA probe preparation, Sheep anti-digoxigenin-POD (Roche Cat# 11633716001) for digoxygenin detection and Clarity Western ECL substrate (Biorad Cat# 1705061) for detection, following the manufacturer’s recommendations.

#### Extension polyA Test (ePAT)

Extension Poly(A) Test (ePAT) was performed adapting (Jänicke *et al*, 2012) protocol. 1 µg of total RNA was denaturated with CS1400 5 min at 80 °C and brought to 37°C 1H with 1 µL of Klenow exo-polymerase (NEB Cat# M0212S) for 3’ extension in 1X First strand buffer, 5mM DTT, 500µM dNTP, and RNAsin(Promega Cat#N2511). Reaction mix was denaturated 5 min at 80°C to inactivate Klenow, and brought to 55°C prior to adding 1 µL of SuperScript III Reverse Transcriptase (Invitrogen Cat# 18080093) incubated 1H at 55°C. The resulting cDNA was amplified using the universal reverse primer (CS1402) and a gene-specific forward primer with Phusion polymerase (ThermoFisher Cat#F530L) according to the manufacturer’s instructions. In parallel, a TVN-PAT reaction was performed using CS1401, a TVN-anchored primer to visualize the amplicon length without poly(A) (A0). Products were loaded on a 2.5% agarose gel in TBE 1X.

#### Degradation kinetic and RNA half-life measurement

Wild-type yeasts and *upf1*Δ strains were transformed with the pCM189-HIS3opt plasmid, and grown as described in Audebert *et al*.(Audebert *et al*, 2024). Degradation kinetic were performed by addition of doxycycline at 10µg/ml (Sigma-Aldrich Cat#D3447) and cell collection at multiple time points (t_0_, t_15_, and t_30_ minutes). Total RNA was extracted by hot phenol/chloroform extraction and precipitated with ammonium acetate and ethanol.

RNAs were fractionated based on their poly(A) tail length as described in the ***RNA Fractionation by Poly(A) tail length*** section, and RNA half-life measurement was performed by RT-qPCR using MFR918 and LA076 for HIS3-opt, GB1659 and GB1733 for Nanoluciferase standard. Half-life from RT-qPCR results were calculated using a nonlinear regression and the exponential decay function e^-(t-l)*k^, as described in Audebert *et al*. (Audebert *et al*, 2024).

#### Illumina sequencing

TruSeq mRNA stranded Library Preparation Kit (Cat#20020594) from Illumina was used according to the manufacturer’s instruction. Single read 75 sequencing was performed on a NextSeq500 flowcell (Illumina Cat#20024906) at the Pasteur Biomics facility.

#### Nanopore Direct RNA sequencing

For library preparation (SQK-RNA002 kit from Oxford Nanopore Technologies), we used 100–500 ng of total RNA or 50 ng of immunoprecipitated RNA (RIP), previously ribodepleted with Oligo d(T)_25_ magnetic beads selection (NEB Cat#S1419S), following the manufacturer’s instructions. Sequencing was performed on MinION or GridION device, using FLO-MIN106D flow cells (R9.4.1) for 72 hours.

### QUANTIFICATION AND STATISTICAL ANALYSIS

#### Illumina sequencing analysis

Raw sequencing data were demultiplexed using bcl2fastq and assessed for quality with FastQC. Adapter sequences were removed using cutadapt. Reads were aligned to the SC288 reference genome (Saccer3) with the STAR aligner. Exon annotation was carried out using the GTF file for *Saccharomyces cerevisiæ* (R64-1-1.104) from ENSEMBL. Read quantification was performed with featureCounts, a function from the Subread package. Subsequent data processing and analysis were conducted using R, and statistical analysis were conducted using the SARTools package with the “shorth” option for Deseq2 normalization^34^.

Table S3, related to Figure 1, shows the normalized count of Illumina sequencing data.

#### Nanopore Direct RNA sequencing analysis

Raw data were basecalled with Guppy basecaller. Reads were mapped using minimap2 and the option “- k14 -ax splice -uf -G 2000 -secondary=no” to the SC288 reference genome Saccer3.fa. Alignements were sorted and indexed using samtools 1.19. PolyA length was determined using Nanopolish v0.13.2 (Workman *et al*, 2019). Subsequently, data were filtered to retain only reads that passed the quality filter “PASS” and reads assigned in doubloons were filtered using a customed R script (10.5281/zenodo.14615891). Subsequent data processing and analysis were conducted using R, and statistical analysis were conducted using the SARTools package with the “shorth” option for Deseq2 normalization (Varet *et al*, 2016).

Table S4, related to Figure 3, shows the normalized count of Nanopore sequencing data.

## Notes

### Competing Interest Statement

The authors have declared no competing interest.

## REFERENCES

Amrani N, Ganesan R, Kervestin S, Mangus DA, Ghosh S & Jacobson A (2004) A faux 3’-UTR promotes aberrant termination and triggers nonsense-mediated mRNA decay. Nature 432: 112–118

Audebert L, Feuerbach F, Zedan M, Schürch AP, Decourty L, Namane A, Permal E, Weis K, Badis G & Saveanu C (2024) RNA degradation triggered by decapping is largely independent of initial deadenylation. The EMBO Journal: 1–29

Beelman CA & Parker R (1995) Degradation of mRNA in eukaryotes. Cell 81: 179–183

Bouveret E, Rigaut G, Shevchenko A, Wilm M & Séraphin B (2000) A Sm-like protein complex that participates in mRNA degradation. The EMBO Journal 19: 1661–1671

Cao D & Parker R (2003) Computational modeling and experimental analysis of nonsense-mediated decay in yeast. Cell 113: 533–545

Celik A, Baker R, He F & Jacobson A (2017) High-resolution profiling of NMD targets in yeast reveals translational fidelity as a basis for substrate selection. RNA 23: 735–748

Celik A, Kervestin S & Jacobson A (2015) NMD: At the crossroads between translation termination and ribosome recycling. Biochimie 114: 2–9

Chen C-YA & Shyu A-B (2011) Mechanisms of deadenylation-dependent decay. WIREs RNA 2: 167–183

Coller J & Parker R (2004) Eukaryotic mRNA Decapping. Annual Review of Biochemistry 73: 861–890

Cosson B, Couturier A, Chabelskaya S, Kiktev D, Inge-Vechtomov S, Philippe M & Zhouravleva G (2002) Poly(A)-Binding Protein Acts in Translation Termination via Eukaryotic Release Factor 3 Interaction and Does Not Influence [PSIϩ] Propagation. MOL CELL BIOL 22

Czaplinski K, Ruiz-Echevarria MJ, Paushkin SV, Han X, Weng Y, Perlick HA, Dietz HC, Ter-Avanesyan MD & Peltz SW (1998) The surveillance complex interacts with the translation release factors to enhance termination and degrade aberrant mRNAs. Genes Dev 12: 1665–1677

Czarnocka-Cieciura A, Poznański J, Turtola M, Tomecki R, Krawczyk PS, Mroczek S, Orzeł W, Saha U, Jensen TH, Dziembowski A, et al (2024) Modeling of mRNA deadenylation rates reveal a complex relationship between mRNA deadenylation and decay. EMBO J

Dehecq M, Decourty L, Namane A, Proux C, Kanaan J, Le Hir H, Jacquier A & Saveanu C (2018) Nonsense-mediated mRNA decay involves two distinct Upf1-bound complexes. EMBO J 37

Eberle AB, Stalder L, Mathys H, Orozco RZ & Mühlemann O (2008) Posttranscriptional Gene Regulation by Spatial Rearrangement of the 3′ Untranslated Region. PLoS Biol 6: e92

Goyer C, Altmann M, Lee HS, Blanc A, Deshmukh M, Woolford JL, Trachsel H & Sonenberg N (1993) TIF4631 and TIF4632: two yeast genes encoding the high-molecular-weight subunits of the cap-binding protein complex (eukaryotic initiation factor 4F) contain an RNA recognition motif-like sequence and carry out an essential function. Mol Cell Biol 13: 4860–4874

Guan Q, Zheng W, Tang S, Liu X, Zinkel RA, Tsui K-W, Yandell BS & Culbertson MR (2006) Impact of Nonsense-Mediated mRNA Decay on the Global Expression Profile of Budding Yeast. PLOS Genetics 2: e203

He F, Li X, Spatrick P, Casillo R, Dong S & Jacobson A (2003) Genome-wide analysis of mRNAs regulated by the nonsense-mediated and 5’ to 3’ mRNA decay pathways in yeast. Mol Cell 12: 1439–1452

He W & Parker R (2001) The Yeast Cytoplasmic LsmI/Pat1p Complex Protects mRNA 3′ Termini From Partial Degradation. Genetics 158: 1445–1455

Heyer EE & Moore MJ (2016) Redefining the Translational Status of 80S Monosomes. Cell 164: 757–769

Hug N, Longman D & Cáceres JF (2016) Mechanism and regulation of the nonsense-mediated decay pathway. Nucleic Acids Res 44: 1483–1495

Hurt JA, Robertson AD & Burge CB (2013) Global analyses of UPF1 binding and function reveal expanded scope of nonsense-mediated mRNA decay. Genome Res 23: 1636–1650

Ibrahim F, Oppelt J, Maragkakis M & Mourelatos Z (2021) TERA-Seq: true end-to-end sequencing of native RNA molecules for transcriptome characterization. Nucleic Acids Res 49: e115

Jänicke A, Vancuylenberg J, Boag PR, Traven A & Beilharz TH (2012) ePAT: a simple method to tag adenylated RNA to measure poly(A)-tail length and other 3’ RACE applications. RNA 18: 1289– 1295

Kervestin S & Jacobson A (2012) NMD: a multifaceted response to premature translational termination. Nature Reviews Molecular Cell Biology 13: 700–712

Kim YK & Maquat LE (2019) UPFront and center in RNA decay: UPF1 in nonsense-mediated mRNA decay and beyond. RNA 25: 407–422

Kurosaki T, Miyoshi K, Myers JR & Maquat LE (2018) Publisher Correction: NMD-degradome sequencing reveals ribosome-bound intermediates with 3’-end non-templated nucleotides. Nat Struct Mol Biol 25: 1059

Lima SA, Chipman LB, Nicholson AL, Chen Y-H, Yee BA, Yeo GW, Coller J & Pasquinelli AE (2017) Short Poly(A) Tails are a Conserved Feature of Highly Expressed Genes. Nat Struct Mol Biol 24: 1057– 1063

Losson R & Lacroute F (1979) Interference of nonsense mutations with eukaryotic messenger RNA stability. Proc Natl Acad Sci U S A 76: 5134–5137

Malabat C, Feuerbach F, Ma L, Saveanu C & Jacquier A (2015) Quality control of transcription start site selection by nonsense-mediated-mRNA decay. Elife 4

Meijer HA, Bushell M, Hill K, Gant TW, Willis AE, Jones P & de Moor CH (2007) A novel method for poly(A) fractionation reveals a large population of mRNAs with a short poly(A) tail in mammalian cells. Nucleic Acids Research 35: e132

Miller C, Schwalb B, Maier K, Schulz D, Dümcke S, Zacher B, Mayer A, Sydow J, Marcinowski L, Dölken L, et al (2011) Dynamic transcriptome analysis measures rates of mRNA synthesis and decay in yeast. Mol Syst Biol 7: 458

Muhlrad D & Parker R (1994) Premature translational termination triggers mRNA decapping. Nature 370: 578–581

Muhlrad D & Parker R (1999) Aberrant mRNAs with extended 3’ UTRs are substrates for rapid degradation by mRNA surveillance. RNA 5: 1299–1307

Nickless A, Bailis JM & You Z (2017) Control of gene expression through the nonsense-mediated RNA decay pathway. Cell & Bioscience 7: 26

Pechmann S & Frydman J (2013) Evolutionary conservation of codon optimality reveals hidden signatures of cotranslational folding. Nat Struct Mol Biol 20: 237–243

Peltz SW, Brown AH & Jacobson A (1993) mRNA destabilization triggered by premature translational termination depends on at least three cis-acting sequence elements and one trans-acting factor. Genes Dev 7: 1737–1754

Roque S, Cerciat M, Gaugué I, Mora L, Floch AG, Zamaroczy M de, Heurgué-Hamard V & Kervestin S (2015) Interaction between the poly(A)-binding protein Pab1 and the eukaryotic release factor eRF3 regulates translation termination but not mRNA decay in Saccharomyces cerevisiae. RNA 21: 124– 134

Schwartz DC & Parker R (2000) mRNA Decapping in Yeast Requires Dissociation of the Cap Binding Protein, Eukaryotic Translation Initiation Factor 4E. Molecular and Cellular Biology 20: 7933–7942

Subtelny AO, Eichhorn SW, Chen GR, Sive H & Bartel DP (2014) Poly(A)-tail profiling reveals an embryonic switch in translational control. Nature 508: 66–71

Tani H, Imamachi N, Salam KA, Mizutani R, Ijiri K, Irie T, Yada T, Suzuki Y & Akimitsu N (2012) Identification of hundreds of novel UPF1 target transcripts by direct determination of whole transcriptome stability. RNA Biol 9: 1370–1379

Tharun S, He W, Mayes AE, Lennertz P, Beggs JD & Parker R (2000) Yeast Sm-like proteins function in mRNA decapping and decay. Nature 404: 515–518

Tharun S & Parker R (2001) Targeting an mRNA for decapping: displacement of translation factors and association of the Lsm1p-7p complex on deadenylated yeast mRNAs. Mol Cell 8: 1075–1083

Tudek A, Krawczyk PS, Mroczek S, Tomecki R, Turtola M, Matylla-Kulińska K, Jensen TH & Dziembowski A (2021) Global view on the metabolism of RNA poly(A) tails in yeast Saccharomyces cerevisiae. Nat Commun 12: 4951

Wang W, Czaplinski K, Rao Y & Peltz SW (2001) The role of Upf proteins in modulating the translation read-through of nonsense-containing transcripts. The EMBO Journal 20: 880–890

Webster MW, Chen Y-H, Stowell JAW, Alhusaini N, Sweet T, Graveley BR, Coller J & Passmore LA (2018) mRNA Deadenylation Is Coupled to Translation Rates by the Differential Activities of Ccr4-Not Nucleases. Molecular Cell 70: 1089–1100.e8

Wiener D, Antebi Y & Schwartz S (2021) Decoupling of degradation from deadenylation reshapes poly(A) tail length in yeast meiosis. Nat Struct Mol Biol 28: 1038–1049

Workman RE, Tang AD, Tang PS, Jain M, Tyson JR, Razaghi R, Zuzarte PC, Gilpatrick T, Payne A, Quick J, et al (2019) Nanopore native RNA sequencing of a human poly(A) transcriptome. Nat Methods 16: 1297–1305

Zünd D, Gruber AR, Zavolan M & Mühlemann O (2013) Translation-dependent displacement of UPF1 from coding sequences causes its enrichment in 3’ UTRs. Nat Struct Mol Biol 20: 936–943

## REFERENCES STAR METHOD

Ghaemmaghami S, Huh W-K, Bower K, Howson RW, Belle A, Dephoure N, O’Shea EK & Weissman JS (2003) Global analysis of protein expression in yeast. Nature 425: 737–741

Namane A & Saveanu C (2022) Composition and Dynamics of Protein ComplexesProtein complexes Measured by Quantitative Mass SpectrometryMass spectrometry of Affinity-Purified Samples. In Yeast Functional Genomics: Methods and Protocols, Devaux F (ed) pp 225–236. New York, NY: Springer US

Tesina P, Heckel E, Cheng J, Fromont-Racine M, Buschauer R, Kater L, Beatrix B, Berninghausen O, Jacquier A, Becker T, et al (2019) Structure of the 80S ribosome–Xrn1 nuclease complex. Nature Structural & Molecular Biology 26: 275–280

Varet H, Brillet-Guéguen L, Coppée J-Y & Dillies M-A (2016) SARTools: A DESeq2- and EdgeR-Based R Pipeline for Comprehensive Differential Analysis of RNA-Seq Data. PLoS ONE 11: e0157022

