## Supplemental Figures S1 to S4 for "Most Upf1-associated mRNAs have short poly(A) tails, lack a premature termination codon and are targeted by NMD"

Figure S1

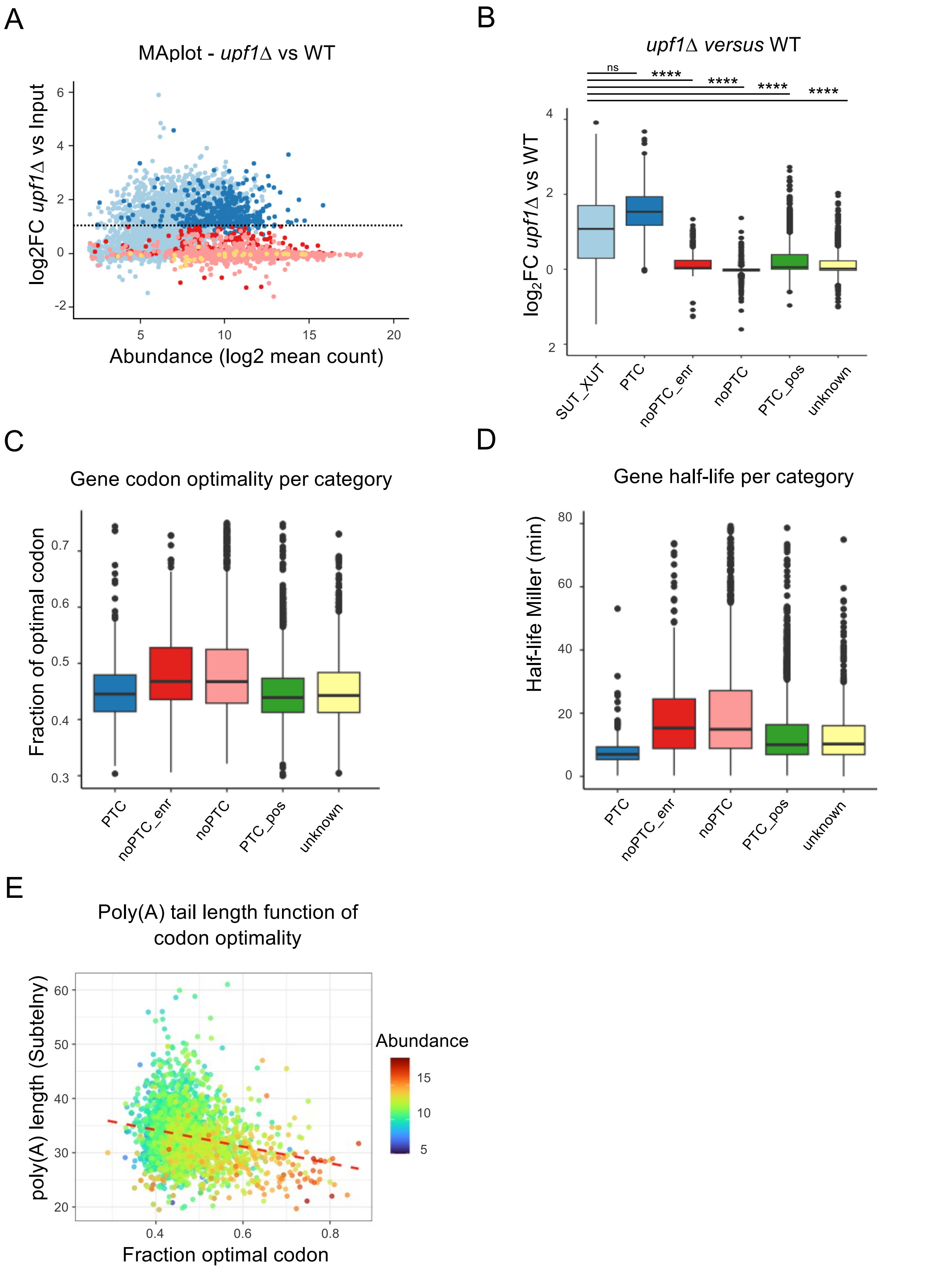
A
MAplot - upf1Δ vs WT
log2FC upf1Δ vs Input
Abundance (log2 mean count)
B
upf1Δ versus WT
ns
****
****
****
****
log2FC upf1Δ vs WT
SUT_XUT
PTC
noPTC_enr
noPTC
PTC_pos
unknown
C
Gene codon optimality per category
Fraction of optimal codon
PTC
noPTC_enr
noPTC
PTC_pos
unknown
D
Gene half-life per category
Half-life Miller (min)
PTC
noPTC_enr
noPTC
PTC_pos
unknown
E
Poly(A) tail length function of codon optimality
poly(A) length (Subtelny)
Fraction optimal codon
Abundance

**Figure S1. Characteristics of transcripts per category, related to Figure 1**

(A) MAplot representing the log2FC between *upf1Δ* and WT as a function of abundance (A = log2 mean count in *upf1Δ* or in Upf1-RIP and WT respectively). Each transcript type is represented as follow: XUT/SUT (light blue), PTC (dark blue), noPTC-Upf1enr (red), noPTC (pink), and mitochondrial transcripts as a control (yellow). Dotted lines indicate a 2-fold threshold (log2FC = 1).
(B) Boxplots comparing *upf1Δ* vs WT log2FC for mentioned categories of transcripts. Pvalues (Tukey HSD test) : ns = p> 0.05, * = p<0,05, ** = p<0,01, *** = p<0,001 and **** = p<0,0001. Detailed pvalues of all Figures are listed Table S2.
(C) Boxplot representing the fraction of optimal codon per gene and per category of transcript, determined according to Pechmann and Frydman, 2013.
(D) Boxplot representing the transcript half-lives (in minute, according to Miller *et al.*, 2011) per category.
(E) Scatter plot representing the relationship between the fraction of optimal codon per gene (abscissa, determined from Pechmann and Frydman, 2013) function of the poly(A) tail length in a WT strain at steady state (ordinate, from Subtelny *et al.*, 2014). Each point represents an individual transcript, colored according to its abundance as indicated by the color gradient. The dashed red line represents the linear regression trend line (Pearson's correlation coefficient, r = -0.219 p value < $2.2\times10^{-16}$).

### Figure S2

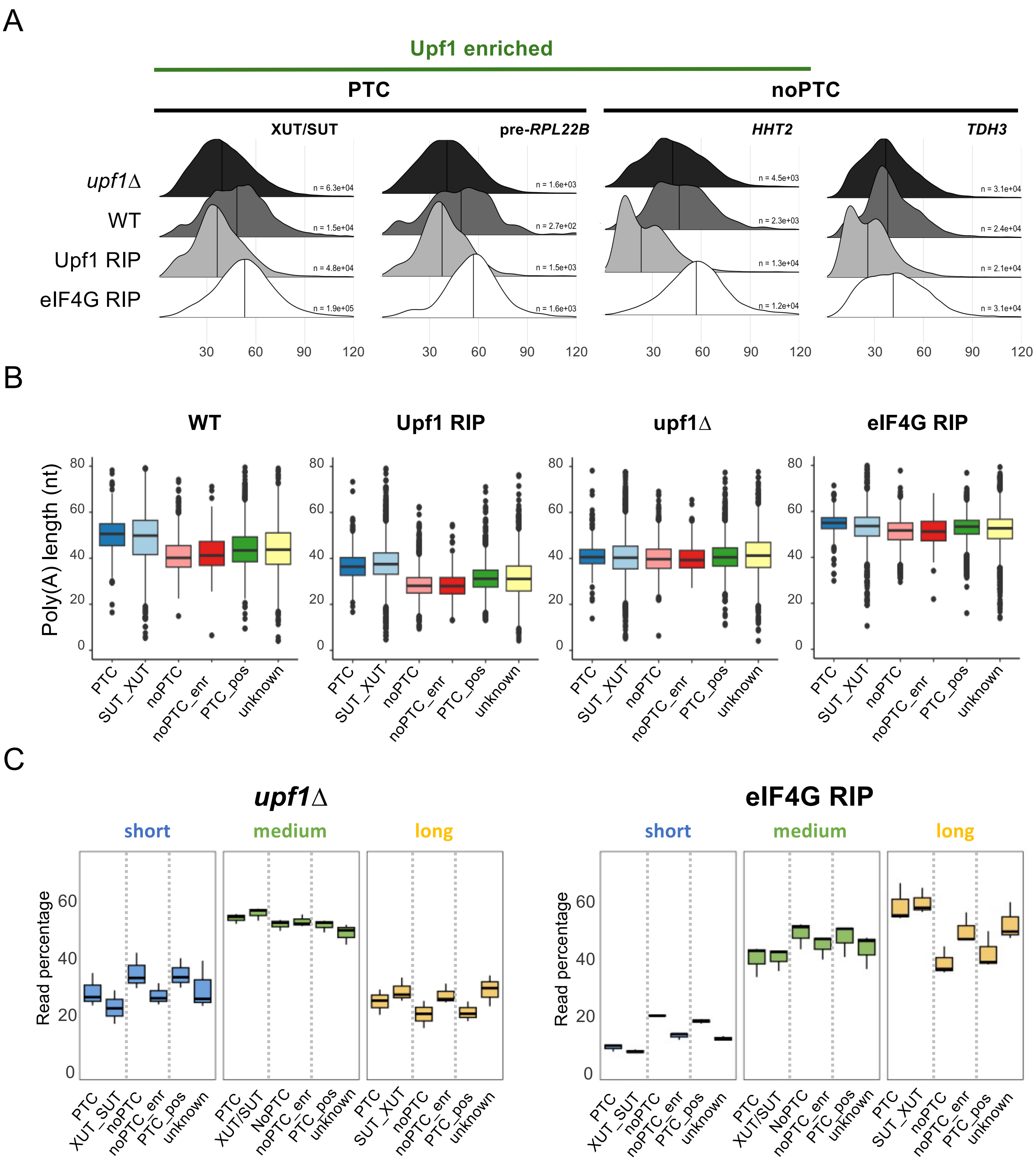
A
Upf1 enriched
PTC
noPTC
XUT/SUT
pre-RPL22B
HHT2
TDH3
upf1Δ
WT
Upf1 RIP
eIF4G RIP
n = 6.3e+04
n = 1.5e+04
n = 4.8e+04
n = 1.9e+05
n = 1.6e+03
n = 2.7e+02
n = 1.5e+03
n = 1.6e+03
n = 4.5e+03
n = 2.3e+03
n = 1.3e+04
n = 1.2e+04
n = 3.1e+04
n = 2.4e+04
n = 2.1e+04
n = 3.1e+04
30 60 90 120
B
WT
Upf1 RIP
upf1Δ
eIF4G RIP
Poly(A) length (nt)
0 20 40 60 80
PTC
SUT_XUT
noPTC
noPTC_enr
PTC_pos
unknown
C
upf1Δ
eIF4G RIP
short
medium
long
Read percentage
0 20 40 60
PTC
XUT_SUT
XUT/SUT
SUT_XUT
noPTC
NoPTC
noPTC_enr
PTC_pos
unknown

**Figure S2. Detailed characteristics of poly(A) tails per category, related to Figure 3**

(A) Ridge plot representing the poly(A) tail length distribution across representative examples in WT (Input) and *upf1Δ* total RNA, as well as Upf1-RIP and eIF4G2-RIP samples. XUT/SUT and *pre-RPL22B* are representative examples of **PTC**-containing NMD targets, HHT2 is a representative example of **noPTC-enr**, and TDH3 is a representative example of the **noPTC** category. Upf1-enriched target are mentioned (green line) while PTC/noPTC categories are indicated in black.
(B) Boxplots showing the proportion of poly(A) tail length in short, medium and long bins across *upf1Δ* and eIF4G-RIP samples. The analysis shows the different sub-categories: PTC (dark blue), SUT/XUT (blue), noPTC (pink), noPTC-enr (red), PTC_poss (green), and unknown (yellow).
(C) Boxplots displaying the proportion of poly(A) tail length in bins (short, medium and long) across *upf1Δ* and eIF4G RIP samples.

Figure S3

A

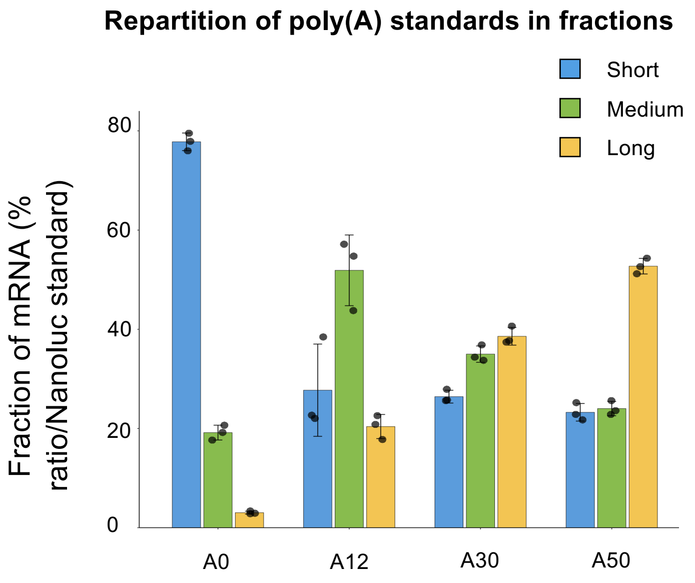

Repartition of poly(A) standards in fractions
Short
Medium
Long
Fraction of mRNA (% ratio/Nanoluc standard)
0
20
40
60
80
A0
A12
A30
A50


B

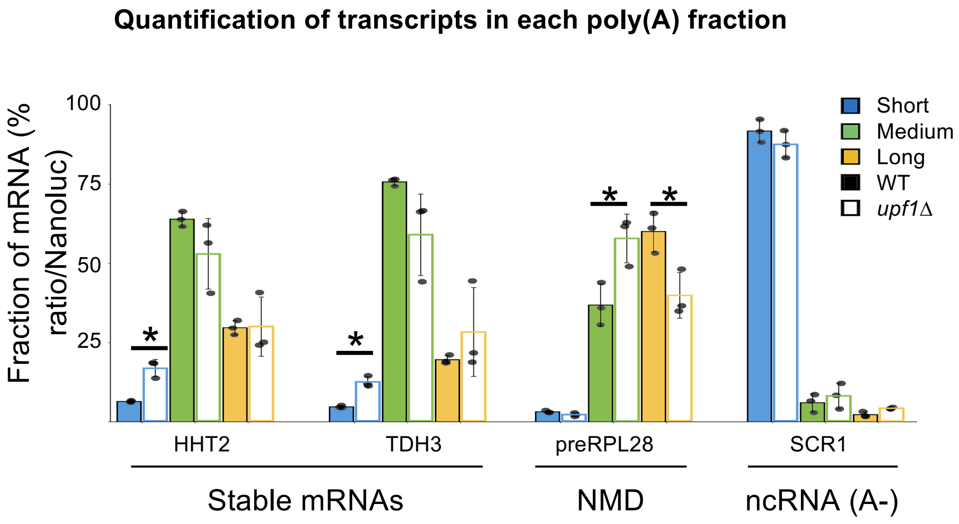

Quantification of transcripts in each poly(A) fraction
Fraction of mRNA (% ratio/Nanoluc)
100
75
50
25
Short
Medium
Long
WT
upf1Δ
*
HHT2
TDH3
preRPL28
SCR1
Stable mRNAs
NMD
ncRNA (A-)

**Figure S3. Relative quantification of poly(A) subfractions during oligodT fractionation, related to Figure 5**

(A) Barplot reporting the relative proportion of each poly(A) tail length in short (S), medium (M), and long (L) fractions of Nanoluciferase mRNA standards with calibrated poly(A) tail lengths (A0, A12, A30 and A50 - see also STAR methods). Data were obtained by RT-qPCR, and show the mean of three biological replicates, black dots represent the value for each replicate, error bar represent the standard deviation (SD).

(B) Barplot reporting the relative proportion of poly(A) tail length in each fractions **short** (blue), **medium** (green) and **long** (yellow) of *ScR1* (deadenylated control), *preRPL28* (NMD control), *HHT2* (non-NMD*) and *TDH3* (non-NMD) transcripts, in wild-type (filled-bar) and *upf1*Δ strains (outlined-bar). Data show the mean of three biological replicates, black dots represent the value for each replicate, error bar represent the SD. * (significant P-values T.test.) : (*preRPL28* – Medium) p = 0.02401; (*preRPL28* – Long) p = 0.02279; (*HHT2* – Short) p = 0.02124; (*TDH3* – Short) p = 0.01165.

### Figure S4

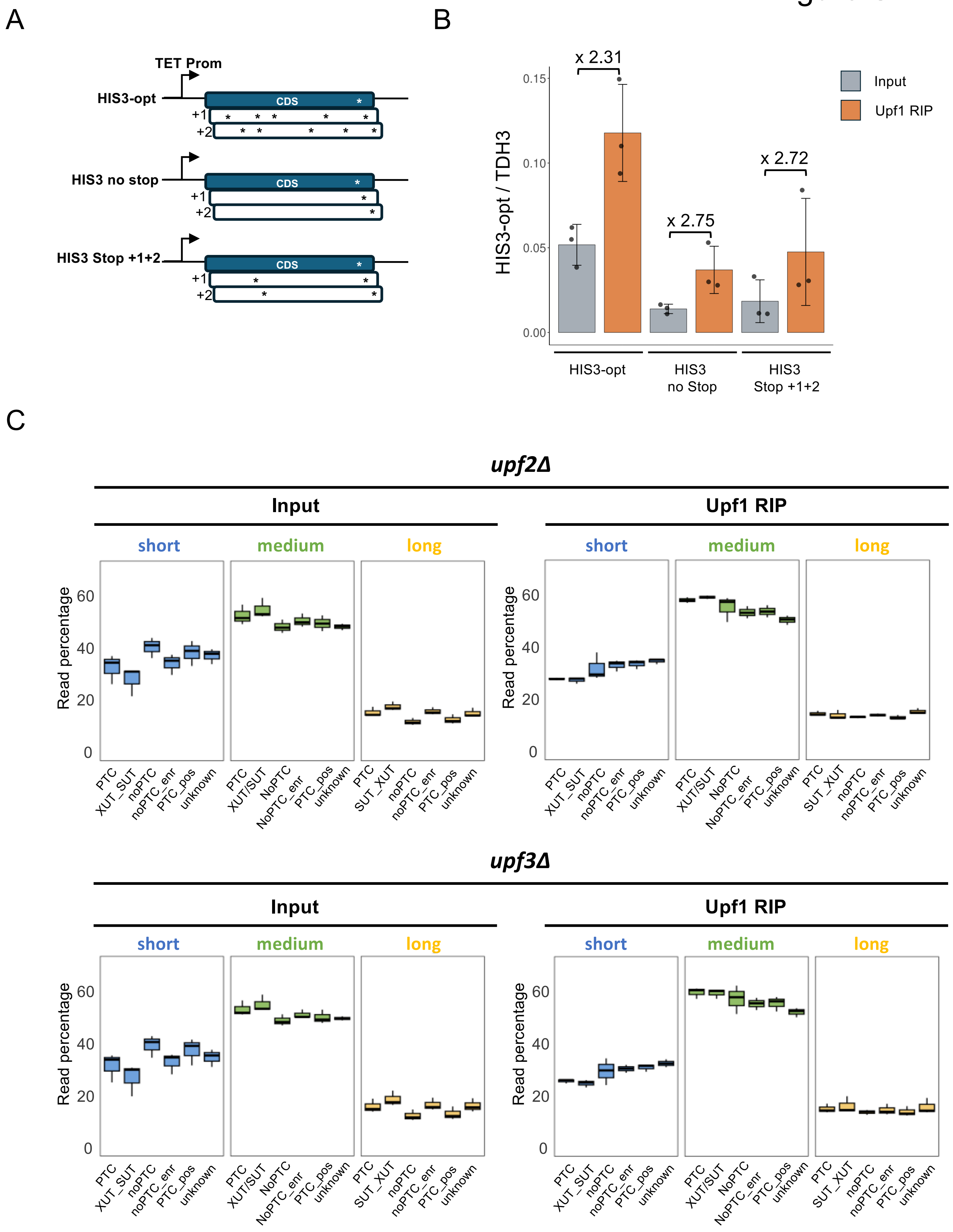

A
TET Prom
HIS3-opt
CDS
+1
+2
HIS3 no stop
CDS
+1
+2
HIS3 Stop +1+2
CDS
+1
+2
B
x 2.31
x 2.75
x 2.72
Input
Upf1 RIP
HIS3-opt / TDH3
0.15
0.10
0.05
0.00
HIS3-opt
HIS3
no Stop
HIS3
Stop +1+2
C
upf2Δ
Input
Upf1 RIP
short
medium
long
Read percentage
60
40
20
0
PTC
XUT_SUT
noPTC
noPTC_enr
PTC_pos
unknown
PTC
XUT/SUT
NoPTC
NoPTC_enr
PTC_pos
unknown
PTC
SUT_XUT
noPTC
noPTC_enr
PTC_pos
unknown
upf3Δ
Input
Upf1 RIP
short
medium
long
Read percentage

**Figure S4. Effect of Upf2 or Upf3 depletion on Upf1-associated transcripts, related to Figure 6**

(A) Schematic representation of the reporters: *HIS3-opt* ("normal"), *HIS3* no stop (no stop in frame +1 or +2), and *HIS3* Stop +1+2 (with a stop in frame +1 and +2). Stop codons are indicated by stars.

(B) Relative quantification of *HIS3-opt* in Upf1-RIP versus WT input, normalized by *TDH3* (non-enriched control). Data show the mean of three biological replicates, black dots represent the value of each replicate, error bar represent the SD.

(C) Boxplot displaying the proportion of poly(A) tail length in bins short, medium and long across Upf1-RIP and Input RNAs for *upf2*Δ or *upf3*Δ strains, comparing indicated transcript categories as in Figure2B.
